# The *C. elegans* Observatory: High-throughput exploration of behavioral aging

**DOI:** 10.1101/2022.06.15.496335

**Authors:** Rex A. Kerr, Antoine Roux, Jerome Goudeau, Cynthia Kenyon

**Affiliations:** Kenyon Lab, Calico Life Sciences LLC, South San Francisco, California, U.S.A.

**Keywords:** aging, automation, behavior, *C. elegans*, lifespan

## Abstract

Organisms undergo a variety of characteristic changes as they age, suggesting a substantial commonality in the mechanistic basis of aging. Experiments in model organisms have revealed a variety of cellular systems that impact lifespan, but technical challenges have prevented a comprehensive evaluation of how these components impact the trajectory of aging, and many components likely remain undiscovered. To facilitate the deeper exploration of aging trajectories at sufficient scale to enable primary screening, we have created the *C. elegans* Observatory, an automated system for monitoring the behavior of group-housed *C. elegans* throughout their lifespans. One Observatory consists of a set of computers running custom software to control an incubator containing custom imaging and motion-control hardware. In its standard configuration, the Observatory cycles through trays of standard 6 cm plates, running four assays per day on up to 576 plates per incubator. High-speed image processing captures a range of behavioral metrics including movement speed and stimulus-induced turning, and a data processing pipeline continuously computes summary statistics. The Observatory software includes a web interface that allows the user to input metadata and to view graphs of the trajectory of behavioral aging as the experiment unfolds. Compared to manual use of a plate-based *C. elegans* tracker, the Observatory reduces effort required by close to two orders of magnitude. Within the Observatory, reducing function of known lifespan genes with RNA interference (RNAi) gives the expected phenotypic changes, including extended motility in *daf-2(RNAi)* and progeria in *hsf-1(RNAi)*. Lifespans scored manually from worms raised in conventional conditions match those scored from images captured by the Observatory. We have used the Observatory for a small candidate-gene screen and identified an extended youthful vigor phenotype for *tank-1(RNAi)* and a progeric phenotype for *cdc-42(RNAi)*. By utilizing the Observatory, it is now feasible to conduct whole-genome screens for an aging-trajectory phenotype, thus greatly increasing our ability to discover and analyze new components of the aging program.

## 1 Introduction

A characteristic decrease over time in capability and health of adults can be observed in the vast majority of species. However, despite its ubiquity, this process of aging has thus far resisted a mechanistic description that accounts for both the nature of the process and its similarity across organisms. Although there are conceptual frameworks that provide an account of how aging could occur—antagonistic pleiotropy, for instance—these give us little insight into the physical basis of the phenomenon. And although a variety of cellular systems have been implicated in regulating aging ([López-Otín et al., 2013]), and many of those are broadly conserved across species ([Taormina et al., 2019]), it is nonetheless unclear how these and, perhaps, other undiscovered systems, cause the aging process to unfold. Thus, the mechanistic basis of aging remains one of the great unsolved mysteries of biology. Additionally, aging has profound practical consequences for human well-being and human society. For instance, in the United States, the cost of medical care for someone in their 70s is more than triple the cost for someone in their 30s ([Dieleman et al., 2016]). Thus, the lack of a mechanistic basis of aging also leaves unrealized one of the areas of greatest cost savings and therapeutic potential in medicine.

Much of what we know about the mechanistic basis of aging comes from experiments in model organisms. Aging experiments are fundamentally challenging because they necessarily take a long period of time, even in comparatively short-lived model organisms. Furthermore, it has historically not been tractable to monitor the full range of phenotypes that might change over an organism’s life. Indeed, the greatest portion of research has used lifespan as a proxy for aging, and for good reasons: death is the ultimate endpoint of aging, the death phenotype is usually unambiguous, and once dead, animals stay dead, thus loosening the requirements for precise temporal control over one’s readout.

However, despite the advantages, there are also sizable drawbacks for using lifespan as a proxy for aging. Firstly, lifespan is low-dimensional: upon discovering a lifespan mutant, although the identity of the gene may tell us about the biological process involved, the lifespan phenotype itself provides no additional clues as to whether the effect is from a known cellular process or an as-yet-unrecognized one. Secondly, critical parts of the mechanism may occur long before death, and therefore the lifespan phenotype may be much less useful in revealing the mechanism than an aging-trajectory phenotype would be. Thirdly, by using lifespan as a primary readout, we leave undetected any processes that affect quality but not quantity of life. For instance, antagonistic pleiotropy anticipates tradeoffs between youthful and aged performance, but monitoring lifespan alone renders us unaware of early-life advantages or detriments.

We therefore sought to examine higher-dimensional phenotypes over time to define an aging trajectory, and to do so at sufficient scale to make practical the discovery of genes involved in biological processes and pathways that are not yet appreciated to have a role in aging. We do not assume such processes necessarily exist, though we suspect that they do; it would be equally informative if, through a reasonably exhaustive search, we could rule out major processes beyond those discovered via lifespan experiments.

*C. elegans* has been a highly useful model organism for aging studies due to a fortuitous combination of factors, including short lifespan, easy genetic manipulation, and a wide variety of genetic and molecular tools. We found behavior to be a particularly appealing readout for quantifying an aging trajectory: different behaviors decline at different ages ([Stein and Murphy, 2012]), monitoring behavior can be noninvasive and minimally perturbative, and a substantial amount of effort has already been devoted to automating measurement of worm behavior ([Husson et al., 2013]), both individually ([Baek et al., 2002, Nagy et al., 2015, Hebert et al., 2021]) or when group-housed ([Ramot et al., 2008, Swierczek et al., 2011, Javer et al., 2018, Pitt et al., 2019]). Furthermore, sufficiently detailed analysis of behaviors has already revealed that genes with common molecular mechanisms tend to cluster together in phenotypic space ([Yemini et al., 2013]). Behavioral analysis has revealed that the EGF pathway acts as a regulator of healthy aging ([Iwasa et al., 2010]) and illuminated a distinction between lifespan and time until cessation of large-scale movement ([Podshivalova et al., 2017]; Oswal et al., unpublished, preprint doi https://doi.org/10.1101/2021.03.31.437415) in *C. elegans*. Although reports of interventions that cause both increased longevity and an improved “healthspan” are common, suggesting a common underlying mechanism, it is unclear to what extent this is due to selection by researchers for more promising phenotypes. A comparison of lifespan and climbing assay performance across the *Drosophila* Genetic Reference Panel showed little if any correlation between the two ([Wilson et al., 2020]), though only one of a variety of age-affected behaviors ([Overman et al., 2022]) was studied. We must surmise that there may be a considerable number of undiscovered regulators of healthy aging that are distinct from known regulators of lifespan.

The search for such regulators would be considerably aided by unbiased whole-genome-scale screening. However, no existing methods appeared to scale sufficiently to allow this at a reasonable level of sensitivity. We therefore sought to create such a method for *C. elegans* behavior. Systems that record many individually-housed animals have the advantage of being able to gather longitudinal data for aging ([Churgin et al., 2017, Pittman et al., 2017]). However, we elected to focus on a system tailored for plate-based group-housed assays, reasoning that the decreased effort to prepare samples and increased throughput was important. Further, we worried that in some formats, longitudinally housed animals might have some of their behaviors governed primarily by interactions with the edge of their necessarily small enclosures, and we wished to have a greater ability to examine behaviors without this being as great a concern.

We therefore decided to extend the Multi-Worm Tracker (MWT) ([Swierczek et al., 2011]), which we had already used to study a variety of behaviors during aging ([Podshivalova et al., 2017]), to operate in a highthroughput automated setting. A particular advantage of the MWT is that it was designed to be highly computationally efficient both in terms of processing time and storage requirements, thereby reducing the danger of computational requirements becoming limiting.

The result of our automation effort is a system that we call the *C. elegans* Observatory. As described below, the *C. elegans* Observatory captures the behavior of worms in a high-throughput automated system, and does so with sufficiently moderate hardware costs and experimenter time such that large-scale screening, such as a whole-genome RNAi screen, is within reach of a single researcher or a small team.

## 2 Materials and Equipment

The development of the *C. elegans* Observatory has taken place over several years, culminating in a system that provides a straightforward workflow for experimentalists (Fig. 1). Through successive rounds of development and refinement, we have continually improved the device, worked around unexpected challenges, and discovered that some strategies were less ideal than we had hoped.

**Figure 1:**
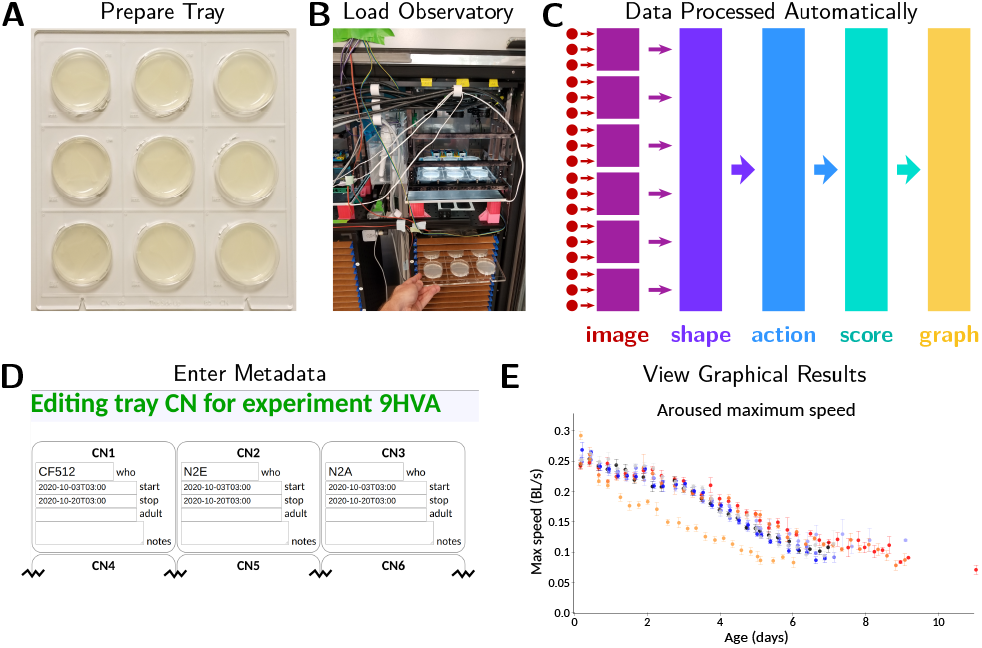
Overview of operating the *C. elegans* Observatory. **(A)** Sample preparation. Worms on standard plates are placed in laser-cut acrylic trays. **(B)** Instrument loading. Trays are placed in a tower in the *C. elegans* Observatory, which automatically cycles the trays for imaging four times a day. **(C)** Data processing pipeline. Triplets of cameras (red dots) send images to six computers (purple squares) for real-time image analysis. Shapes and positions are extracted and sent to a central server (indigo); from these, successive processing steps extract timecourses of behavioral parameters (light blue), per-animal health scores (teal), and finally graphs of behavior over time (gold). All analysis happens automatically with no user intervention necessary. **(D)** Metadata entry. Users enter metadata identifying what sample is on each plate. **(E)** Online graphical results. Metadata is used to group samples and present graphs to users on request, as data becomes available.

Here, we describe the critical pieces of hardware and the design principles that have proved important, pointing out where appropriate both what we actually did during development and what a superior solution would be if building another system. The descriptions are not intended to be adequate to create a part-for-part identical copy, nor would this be advisable. Indeed, it would be a substantial engineering effort for us to create a duplicate system ourselves. However, as we continue to work on the system, we will make any parts lists and assembly diagrams that we generate available at https://github.com/calico/elegans-observatory, in addition to the latest versions of the source code, with the ultimate goal of fully documenting the necessary steps to create additional instances of the Observatory. If other groups are interested in reproducing the Observatory, we are happy to provide guidance and to accelerate completion of the public repository.

### 2.1 Observatory Hardware

Physically, the Observatory consists of an incubator with motion-control, sample holding, and imaging systems, plus an associated computer tower (Fig. 2, central panel). Subcomponents are described below.

**Figure 2:**
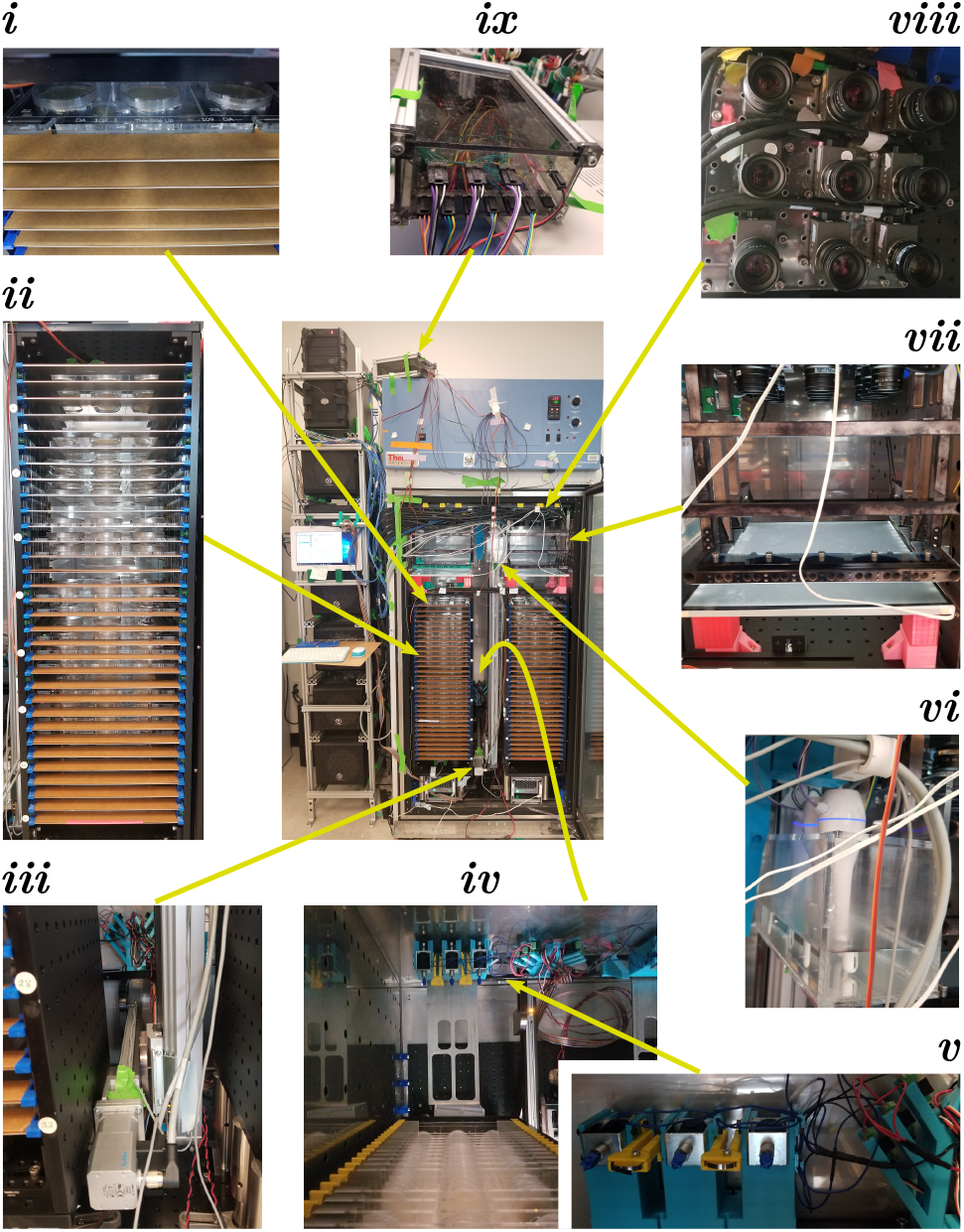
*C. elegans* Observatory Hardware. Center: Assembled Observatory including computer tower (left) and incubator with custom hardware (right). Worms on 6 cm plates are mounted in acrylic trays that are placed into a tower (*i*); the full tower contains 32 slots for trays (*ii*), and there are two towers per incubator. Trays are moved via a two-axis linear motion system (*iii*), with a forklift-style assembly to lift the trays from behind (*iv*). The forklift also braces the tray in position when on the imaging platform and contains solenoids to deliver mechanical stimuli (*v*). A custom humidifier box (tank + mini desktop humidifiers) maintains humidity (*vi*). The imaging platform (*vii*) consists of a light box mounted on top of the tower and a frame mounted to the ceiling that holds the tray in fixed position relative to the ceiling-mounted camera array (*viii*). Temperature and humidity are monitored and maintained with the aid of a custom electronics box (*ix*) containing a small arduino-style processor (Teensy 3.2) that reads sensors and controls humidifiers and the incubator chiller.

#### 2.1.1 Incubator and Environmental Control

Because *C. elegans* lifespan is strongly temperature-dependent and we have a variety of heat sources within the incubator, we use a Thermo Scientific Forma 5920 incubator with lateral air flow, as in ([Stroustrup et al., 2013]), as the path and volume of airflow reduces temperature gradients compared to other designs. To provide sufficient attachment points for hardware, we mounted 75 × 45 cm ThorLabs breadboards (MB4575/M) to the floor and ceiling using custom 3D-printed brackets that hook into holes in the incubator walls. The 3D printing orientation was chosen for maximum strength to sheer forces applied vertically; both ABS and PLA were used as materials.

To reduce mold and bacterial contamination, we placed two small air purifiers on the floor of the unit. During the time of construction, HEPA purifiers of appropriate size were unavailable at our location, so instead we used electrostatic plate air purifiers (Alford Industries HexaOne—now apparently discontinued, but as of this writing the same unit is available under the brand name Nectar HexaOne or DWMHexaOne).

Heating caused by the electronics in the incubator would have caused the internal temperature to exceed 25 °C had we relied only on passive cooling. Unfortunately, the aggressive chiller/heater logic used in the Forma 5920 incubator, though good for tight and rapid temperature control, resulted in low (∼ 40%) relative humidity which caused excessive drying of worm plates. We therefore spliced into the chiller power control line (equivalent to flipping the incubator’s front panel “refrigeration” switch), selected a target set point lower than the desired temperature, and wrote our own logic to run the chiller as needed. A cycle time of 90 seconds was chosen as a compromise between temperature stability and possible wear-and-tear on the chiller. Under normal conditions, the chiller is on for approximately 30 seconds out of every 90, which is adequate to maintain a stable temperature of 24.6 ± 0.2 °C. Note that the desired temperature can be adjusted in software; we have also run experiments at 20 °C and 15 °C. However, for this study, the temperature-sensitive sterile strain we used required 25 °C.

Although our modifications reduced the rate of dehumidification, the resting humidity level was still too low when only a few plates were in the incubator. To raise the relative humidity to ∼ 80%, we built a box with capacity to hold roughly one liter of water; the box was constructed from laser-cut panels of 6 mm thickness acrylic glued together with acrylic cement (IPS corporation #10315, #16 fast set, clear, medium bodied solvent cement). We then placed three desktop humidifiers (Jaywayne Portable Mini USB Humidifier) into this box (Figure 2 detail *vi*) under programmatic control via splicing into their on/off switches.

Although this scheme was effective at maintaining humidity and temperature in the desired range, we recommend an incubator with humidity control instead.

#### 2.1.2 *C. elegans* Housing

The *C. elegans* Observatory is designed to use standard 6 cm worm plates for the convenience of experimenters. Because precise dimensions differ between manufacturers, we had to settle on a particular model; we chose Fisher Brand FB0875713A. We laser cut 25 × 26 cm trays out of 6 mm thick acrylic to hold nine plates at a time (Fig. 1A), using a precisely-sized circular socket (hole) to allow plates to be inserted and held firmly without a clamp mechanism. The design we used, viewable in the files used for laser cutting in https://doi.org/10.5281/zenodo.6645842, is effective but could be improved further; the thin elements tend to break, albeit without compromising adequate function. Additionally, we engraved human-readable and machinereadable identifiers by every plate socket so the identity of each position is unambiguous merely from viewing an image. This prevents any misattribution, providing that the user correctly provides metadata about the strain and condition at a particular position.

To enable precise alignment of the tray, we cut a pair of triangular notches, with tips cut out in circles (visible at the bottom of Fig. 1A), that slide into matching triangular teeth in a brace on the imaging platform. This allows repeatability of a few tens of microns between successive deliveries of the tray. Trays are named via a two-letter code with a decimal decimal equivalent for redundancy; these were engraved at the bottom of each tray (visible in Fig. 1A). Because mistaking the tray identity would be a critical error, the user interface requires experimenters to enter both the letter-code and the numeric code in metadata; it alerts the user if there is a mismatch.

To store the trays in the incubator, we built two tray towers that each hold 32 trays on C-shaped sheet metal shelves with 25 mm vertical spacing (Fig. 2 detail *ii*). We also tried 20 mm spacing, which was workable in principle but in practice it was challenging to meet positional targets with sufficient accuracy, so we settled on the wider spacing. The open part of the C faces backward to allow the forklift to lift the tray from the center (Fig. 2 detail *iv*). The outside walls of the tower are made from 90 × 30 cm breadboards (ThorLabs MB3090/M) for the sides and 30 × 30 cm breadboards for floor and ceiling (ThorLabs MB3030/M). The breadboards are attached in a box shape via metal or 3D-printed plastic angled brackets, with the side walls inset to be flush with the edges of the top and bottom breadboards. Within this open box, oriented with the long axis vertical, we attached sheet metal shelves (dimensions 26 × 30 cm, 4 cm wide rim along the long edges and one short edge) via a combination of 3D-printed retaining brackets (blue in front, visible in Fig. 2 detail *ii*; yellow in back, visible in Fig. 2 detail *iv*) and laser-cut 6 mm acrylic guide rails (oriented vertically, attached to the inner sides of the box via 3D-printed brackets, with notches to allow the shelves to be inserted). A superior design would have been to include small notches on the metal shelves to allow the shelves to be retained by an internal element instead of the yellow back retainers.

Additional features of the towers include optical posts for legs (Thorlabs P150/M), allowing easier removal than if the box were affixed to the incubator’s floor breadboard, and guide strips of 1 mm thick acrylic with triangular alignment teeth to allow precise hand-loading of trays by drawing the corresponding triangular notches in the trays against the guide teeth. We left the paper coating on one side of the acrylic to aid visual alignment (Fig. 2 detail *i*).

#### 2.1.3 Imaging Platform

Successful imaging requires high-quality illumination. We judged that the spatial constraints in the incubator favored brightfield imaging, where optical elements can be stacked vertically, rather than darkfield where illumination sources must be placed at an oblique angle. Because *C. elegans* in brightfield generates much of its contrast by scattering rather than absorption, it is important that the incident light be reasonably collimated. To achieve a broad and uniform collimated light field, we used a custom 12^”^ × 12^”^ LED light panel (GLLS, Green LED Lighting Solutions; 5700K color temperature) covered by a diffusing sheet (Lee Filters #216) to increase uniformity, followed by a pair of computer privacy screens (we tested several vendors’ and all worked fine) cut to size and placed at 90° angles. We then designed a metal housing, affixed to the ceiling, to hold trays roughly 5 cm above the light panel (Fig. 2 detail *vii*). The housing consists of mostly open side walls connected with braces in the front but open in the back; attachment points were machined into the side walls to allow 3D-printed guide and retaining wedges for precise tray alignment. Final position was achieved by pushing the tray into place against a 6 mm thick laser-cut acrylic guide bar with two triangular alignment teeth (visible on top of the lowest brace in Fig. 2 detail *vii*; alignment notch visible at the bottom of the tray in Fig. 1A).

This design allowed highly reproducible positioning (*<* 40 µm) and acceptable contrast (*>* 1.6 : 1 between background and worm body).

#### 2.1.4 Motion and Stimulus Delivery

The core of the motion system consists of a pair of Festo linear stages (Fig. 2 detail *iii*): a larger vertical stage (EGC-80-1200-TB-KF-0H-GKZUB-6M2C with motor EMMS-ST-57-M-SEB-G2 and gear unit EMGA-60-P-G5-SST-57) clamped to both floor and ceiling breadboards in the incubator, and a smaller horizontal stage (EGSK-33-300-6P with motor EMMS-ST-42-S-SE-G2) mounted on the vertical one and on which the forklift tines are mounted. These run a fixed pattern of motions generated via a program written in CODESYS 2.3 (Festo variant) running on a dedicated NUC-style mini-PC using the Microsoft Windows 10 operating system. We had judged that the increased reliability of a stand-alone motion system was important. However, given the high in-practice reliability of our server—higher than the motion control system—this concern proved unwarranted, and allowing the server to coordinate motion would have been the superior option.

To physically move the trays, a horizontal bar was placed at right angles to the horizontal stage (Fig. 2 detail *iv*). On either side of this bar, aligned with the left and right towers, we attached a broad single-tine forklift blade (Fig. 2 detail *iv*) with a beveled front edge to reduce the chance of colliding with a tray (not shown). Above the blade we also mounted a 3D-printed bracing assembly with circular coils of spring wire (Fig. 2 detail *iv*; silver band in front of yellow mounting arm; two layers of 301 spring tempered stainless steel, 0.005^”^ × 1*/*8^”^, Lyon Industries) to contact the tray with appropriate force.

In order to help determine animals’ capability rather than their intrinsic drive, we wished to provide an aversive stimulus to induce activity. We selected mechanical tap as an easy and precisely-deliverable stimulus. We therefore mounted solenoids (RobotGeek ASM-SOL-MD) aligned with the spring-wire brace (Fig. 2 detail *iv*; silver boxes with blue plastic-tipped probes).

Mechanical stimuli are triggered as part of the consensus-detection algorithm that also signals the initiation of a recording session. The striking of the solenoids are controlled by custom real-time stimulus delivery and voltage monitoring software running on an Arduino-compatible Teensy 3.2 computer-on-a-chip (PJRC). High-speed imaging confirmed that the impacts provide sufficient acceleration at all positions on the tray to expect a tap withdrawal response (data not shown), and observation of animals confirmed this. Unfortunately, the mechanical design of the system admitted longer-lasting vibration than previous non-automated rigs, reducing the reliability of our existing algorithms to detect tap-induced behavior such as reversals. We therefore focused on highly reliable general activity measures (speed) pending either our development of high-reliability response-detection algorithms despite vibration, or improved vibration control, or both.

#### 2.1.5 Camera Array

To image the plates of worms, we installed two 3 × 3 arrays of 5 megapixel USB3 cameras (Pixelink PL-D725MU-T, 2592 × 2048 8-bit grayscale, USB-3 interface, maximum frame rate 75 Hz) with 25 mm focal length lenses (Navitar NMV-25M1) situated with the camera sensors roughly 25 cm above the positions of the plates in trays mounted in the left and right imaging platforms (Fig. 2 detail *viii*). This provided an appropriate field of view with one pixel corresponding to a 40 µm square. We ran the cameras at 50 frames per second with 19 ms exposure.

The magnification achieved with the cameras, which yields images with 40 µm/pixel, is moderately less than in previous studies using the MWT. If a more precise estimate of worm posture were desired, higher resolution cameras could be chosen to yield equal (29 µm/pixel from a 12 megapixel sensor) or better (20 µm/pixel from a 20 megapixel sensor) resolving power as compared to previous studies. This would, however, necessitate some decrease in frame rate unless a 10 GigE interface were used instead of USB-3.

To affix the cameras to the ceiling breadboard, we used a laser-cut acrylic mounting sheet with camera positions predefined. Because of the low rigidity of the acrylic, it was necessary to cut fixed holes to allow screws to be placed into the breadboard near to the cameras. This solution was chosen for expediency as we were still adjusting the tray format at the time. With the tray format now defined, the superior solution would be a machined aluminum mounting plate to allow all cameras to be adjusted together and affixed with clamps at the edge of the mounting plate.

Residual discrepancies between the precise centering of each camera over each plate were corrected in software. Focus and aperture were adjusted by hand to give a sharp image and bright but non-saturated background when looking at a plate (8-bit pixel values of 200 or higher).

#### 2.1.6 Environmental Monitoring and Control

In order to monitor the temperature and humidity inside the incubator, and verify that there were no variations in temperature and humidity large enough to confound our results, we installed six temperature/humidity sensors (DHT22; Adafruit #Ada385) and modified the Ticklish software to read the temperature/humidity values using a Teensy 3.2 device. We calibrated the sensors’ temperature readings by placing them together with an alcohol lab thermometer; we assumed the sensors’ mean humidity reading was correct. Each sensor’s readout was then corrected in software to match the ground truth result. The sensors were then deployed in a variety of locations throughout the incubator. Unfortunately, the DHT22 sensors have not proven very reliable under the environmental conditions in the incubator, and we have had to replace them multiple times. A more robust solution would be preferable.

We used the Teensy 3.2 device to run the desktop humidifiers and the incubator’s chiller as well as to read the DHT22 sensors. The Teensy 3.2 chip, together with wiring and the minimal circuitry needed, were placed in a custom box (Figure 2 detail *ix*) made from aluminum rails and lasercut acrylic.

### 2.2 Observatory Computational Resources

In order to acquire images from the 18 cameras, we custom-built six PCs (“imaging computers”) with Intel i7-6700 CPUs, 16 GB of RAM, a 512 GB local SSD, and a PCIe four-port full speed USB3 expansion card (Renesas, though we have used other vendors also). For operating system, we installed Ubuntu 16.04 LTS. Because the Multi-Worm Tracker software is highly efficient, this provided more than adequate computation to track and segment the animals from three cameras simultaneously. The fourth USB port is unused. To avoid possible breakage or intrusion, we keep these computers disconnected from the internet and do not apply any updates.

To gather the results and coordinate the separate computers, we also built a custom server PC that included the following components: Intel i7-6700K CPU, 32 GB RAM, 128 GB SSD boot drive, 512 GB NVMe M.2 SSD data transfer drive, 14 TB raw data storage hard drive (“Tier 1” pipeline output), 6 TB analyzed data storage hard drive (other tiers), and 6 TB auxiliary hard drive. Again, we used Ubuntu 16.04 LTS as the operating system. The server has both a connection to our internal network and, through a second NIC and a switch (NetGear ProSafe 8 port GB ethernet switch), a local network shared only with the imaging computers. The server PC shares via NFS one subdirectory on the data transfer drive with each of the imaging computers. The imaging computers use this to transfer data to the server and to coordinate activity. The server PC also mounts our cluster filesystem, allowing backup of all data as well as the ability to serve the user interface from a cluster node.

Although we elected to use our local cluster filesystem, using cloud-based storage should also be possible. We chose the local approach mostly for ease of use and speed of access for operations like traversing the entire directory structure. There is no reason in principle that cloud storage could not be used; one would just have to be thoughtful about caching certain types of information to avoid excessive access. Additionally, one might need a more vigorous security policy if the server had greater exposure to the outside world.

The motion control software required the Windows operating system, for which we used a NUC-style mini PC (smaller than an actual Intel NUC, though in the future we would just use an Intel NUC). For stability, this also is disconnected from all networks and operated without updates. Of the computers, only the last requires a keyboard, mouse, and monitor. The others are accessed via the network: the server is accessed directly, while the imaging PCs can only be accessed through the server.

## 3 Methods

### 3.1 Hardware Manufacturing and Assembly

Because the instrument was designed and built over a period of years, the assembly process was highly ad-hoc. We followed several principles during the design and construction process. First, if in-house 3D printing or laser-cut acrylic could suffice for a mechanical component, we used that rather than commercial parts or custom machining. Second, when laser-cut acrylic would suffice, we would use that due to the greater speed and accuracy, and mostly superior mechanical properties, as opposed to additive 3D printing. Third, when making 3D-printed parts, we paid special attention to the layer adhesion of the material, as this is typically critical in parts that are load bearing along multiple axes; we found PLA-based materials worked fairly well from among the materials we tried, though as manufacturers introduced new products we found that we were best informed if we tried each material. Fourth, tapping holes (i.e. cutting in screw threads with the appropriate cutting tool) in either acrylic or 3D-printed materials is fast and easy, so we took extensive advantage of this.

Most 3D-printed parts were printed on a LulzBot Mini 1.0. A variety of 40W-60W laser cutters were used to cut acrylic. Sheet metal cutting and machining were outsourced. 3D parts were designed with SolidWorks 16 (Dassault Systèmes), OpenSCAD 2015.03 or later, and/or Onshape. 2D parts were designed with SolidWorks 16, manually in Inkscape 0.92, or programmatically (written to SVG files).

Assembly of the instrument proceeded broadly in this order: first, breadboards were installed on the floor and ceiling of the incubator. Second, the vertical stage was installed by using 3D-printed mounting blocks to hold the stage and fill the excess space between floor and ceiling. Third, the horizontal stage and forklift assembly was installed. Fourth, we added the housing to hold trays above the illumination platform, as the housing is designed to screw directly into the breadboard and thus isn’t very movable. Fifth, the towers were situated under the tray-holding housing. Sixth, the light source was added on top of the tower. Seventh, the cameras were installed on the ceiling (unscrewing the front braces from the imaging frame to allow easier access). Finally, all remaining components were added (sensors, humidifier, etc.).

A custom aluminum-frame rack was constructed next to the incubator for the imaging and server computers, which were situated in the rack with 3D-printed mounting brackets. The ethernet switches, motor controllers, and such, were also attached to this rack via 3D-printed brackets.

### 3.2 Software

#### 3.2.1 Programming Languages

Our philosophy has been to select languages that are particularly appropriate for the most challenging computational tasks that we face, rather than to pick a single language and then solve challenges to the extent possible within the constrains of that language. Image processing and hardware control was therefore written in C++ (compiled with GCC 7.5) for its performance and low-level hardware access, with one module in Rust (version 1.52 or later) due to Rust’s equally good performance and superior facilities for handling complex data structures. Post-capture analysis was mostly written in Scala, as it has excellent (Java-class) performance and JVM compatibility, and its type system prevents broad classes of bugs thereby increasing stability despite a development team consisting primarily of a single person. Some older post-capture code (Choreography) was written in Java, and one part was written in Rust for additional speed and confidence in correctness. Motion control was written in CoDeSys 2.3, and some coordination tasks were written in Bourne Again SHell (bash, version 4.3).

Although the benefits of this approach have been considerable, it is worth noting one major drawback: few people have the requisite skills to be able to rapidly begin development on any arbitrary part of the system. Indeed, we were not familiar with all the tools when we began to use them. Thus, although the benefits we gained will also apply to anyone else—for instance, the Scala and Rust type systems prevent broad classes of errors common to new developers, and to the original developer who has not looked at the code in too long and forgotten some essential details—the barrier to entry is higher than for the typical project written in Python, R, or Matlab. In our defense, we can only add that mastering the tools used will also considerably advance one’s skill as a programmer via exposure to the concepts and strategies common to the different languages.

#### 3.2.2 Motion Control

The motor controllers (Festo CMMO-ST-C5-1-DION) for the stages were attached, as recommended, to a programmable logic controller (PLC), Festo CPX-GE-EV-S. We used the provided CoDeSys 2.3 environment to develop a control program for the PLC, based around preprogrammed locations and step sizes downloaded into the motor controllers. We used mostly the continuous function chart (CFC) programming language, which represents data flow as occurring through wires between control blocks; we first developed primitives for tasks such as picking up a tray from tower, chained these primitives together to cycle through all the trays, and chained these cycles together to make a multiday protocol that would run independently of external control. In retrospect, using a serial-over-USB or similar controller, commanded from our server via a program written in a standard procedural or functional programming language, would have simplified all aspects of the process with equal or greater reliability.

#### 3.2.3 Ticklish

In order to deliver precisely-timed stimuli, it is helpful to have dedicated hardware that is not subject to the same type of operating-system induced delays that a general purpose computer is. Although one can purchase hardware specifically for signal generation, we elected instead to write custom software for the Arduino-compatible Teensy 3.2 processor on a chip. We defined a set of commands to be delivered via serial-over-USB to the Arduino device, specifying a protocol of precisely timed (typical temporal precision < 100 µs) voltage switches on desired output pins. We also allowed live commands to query status including reading voltage on pins. Once a protocol is downloaded, the device then runs it independently of the host computer, preventing any asynchronous interrupts or heavy workloads from interfering with precise timing. Our temperature/humidity sensors offered a slightly more complex trigger-and-readout scheme, which we also implemented in the software, though this readout is somewhat slow and therefore could delay a protocol that is running at at the same time. However, as we use two separate Teensy 3.2 devices to deliver taps and to sense and control the environment, and the latter does not require precise timing, the readout delay is not an issue in practice.

Note that precise timing is important for the operation of the solenoid-based tappers. Typically, only a few tens of milliseconds of current are sufficient to drive the solenoid to collide with the tray; therefore, relatively small (millisecond-scale) variations can change the nature of the tap from a ballistic-like impact to a strike-and-briefly-hold motion.

We named the software Ticklish as our primary use for it is to deliver mechanical stimuli. Because it works as a general-purpose stimulus delivery and voltage querying device, we gave it its own repository (https://github.com/Ichoran/ticklish). It is written in Arduino-themed C++.

#### 3.2.4 The Multi-Worm Tracker

The detection and segmentation of worms from background is accomplished via the Multi-Worm Tracker ([Swierczek et al., 2011]). We separated the core image processing routines from the LabView-based UI and established a new repository for maintenance and development of the image processing portion (https://github.com/Ichoran/mwt-core). The core routines remain essentially unchanged. In brief, the MWT calculates a decaying average of the image to estimate a background, then segments worms as differences from background that are over threshold. In order to better detect moving worms, we amended the decay algorithm to be asymmetric: to quickly accept darkening pixels but only slowly lighten them. This better detects slower-moving animals that gradually move into a new area. Regions that pass a threshold are flood-filled using a less stringent threshold and the outer contour is extracted and saved. The region around detected animals no longer updates its background, so a once-moving animal stays at a high contrast from the local background even if it stops moving.

We also added SIMD-based commands to project an image onto the horizontal or vertical axis; these are used to help detect the octicon symbol block to identify trays and to detect the position of high-contrast edges to help compensate for vibration. The operation is trivial (summing pixels along rows or columns); only the use of SIMD instructions (with a roughly 4x speedup, though we did not measure carefully) is noteworthy.

#### 3.2.5 Tray Identification

Because the motion automation is decoupled from both the imaging and server computers, yet the arrival of a tray requires a prompt and synchronized response of these computers, the robust and rapid detection of the tray is essential. We created a custom numbering scheme for trays and a custom set of eight easily line-engraved characters, “octicons”, to read via template matching; by engraving both the human-readable and this machine-readable format next to every slot in every tray, we were able to rapidly and uniquely identify every slot both computationally and visually. More information about the implementation is available in the Supplementary Materials and in Supplementary Figure 1.

#### 3.2.6 Spanner

The central task of the imaging computers is to detect when a tray has arrived and to run the MWT to segment worms when it has. This task is made more complex because of the need to establish two types of consensus. First, one imaging computer sees only one column of three plates, not the whole tray, so the start of any protocol has to be made in coordination with the other imaging computers who may have a different timeline for positive identification of the tray. Secondly, within the imaging computer itself, the identification from the three different image streams has to be coordinated.

To perform these operations, we wrote custom software, Spanner. Spanner is written mostly in C++ (C++14 dialect); it capture frames from the camera using vendor-supplied Pixelink camera drivers, detects when an octicon block appears in a per-camera specified region, and determines a consensus detection time across the three cameras it controls. In order to include adequate handling of error conditions, we developed a state machine of nontrivial size to seek the octicon block, positively identify the tray for a predefined amount of time (to ensure stability had been achieved—the system observes the tray in motion before it reaches its final braced position), and coordinate across cameras. We wrote the state machine logic module in Rust (2018 dialect) instead of C++, as its pattern-matching capacity simplifies the logic of state switching while also considerably reducing the chance of error. For details, one should refer to the source code.

Additionally, Spanner saves a snapshot image at the beginning and end of each time window where we have decided to measure speeds. Thus, the complete data consists of six full-frame images plus a timecourse of outer contour and centroid position for each detected moving object. Note that due to collisions, one animal may be found as distinct temporally separated objects.

#### 3.2.7 Controller

A key function of the server computer is to coordinate the different imaging computers so that a behavioral protocol is run in a way that is synchronized across those computers. Another key function of the server is to monitor and control environmental parameters (temperature and humidity). Because both of these functions require attention to timing and communication with Teensy devices running Ticklish, we decided to combine the functions into a single program, written in Scala, which we called Controller.

The behavioral protocol aspect of Controller functions as a simple synchrony-detection device: it watches a set of directories for the appearance of a file to indicate a tap request; if enough (in practice, two) requests come in within one second, the commands to run the standard behavioral protocol are sent to the Teensy device connected to the solenoids that impact the trays. It also writes a file in each directory that specifies the time at which the protocol will start. Each imaging computer (via Spanner) reads this file when it appears and waits until the specified time to begin processing the image stream. The server computer functions as an NTP server for the imaging computers, which in practice generally results in time synchrony of under one frame (20 ms).

The environmental control runs on simple threshold-based logic; if heat is too high or humidity too low (based on the average value of the sensors, with any nonworking sensors ignored), the chiller or humidifiers, respectively, are turned on; how long they are turned on depends on how many thresholds are exceeded. This is very simple to tune, though it provides less precision and speed of return to baseline than would a software-based PID controller with appropriate parameters. The environmental control portion also outputs the environmental parameters to a log file, which is then conveyed to the web interface.

#### 3.2.8 Pipeline

The Spanner software extracts the outlines and centroids of worms in each frame and stores them locally. However, this leaves a sizable amount of processing necessary to obtain any biological insights: the data is distributed, is organized by camera instead of sample identity, and is very low-level. To assist with rapid development of insight, we built a processing pipeline that runs in batch mode after each recording from a tray (Fig. 3A) and computes summary data as the data becomes available.

**Figure 3:**
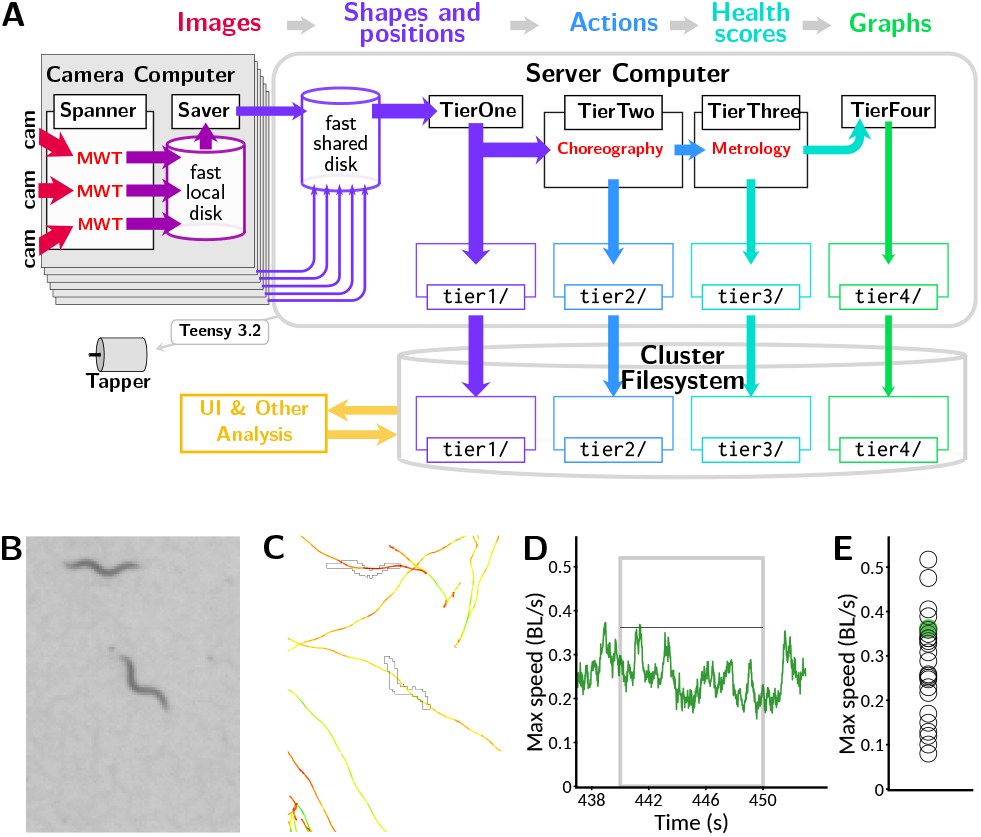
Data processing pipeline and data reduction. **(A)** Information flow within pipeline. Image data from cameras (red: 2.3 TB per session across 18 plates imaged simultaneously) is processed in real time with the MWT software into animal shapes and positions (purple: 1 GB per session). This is transmitted to a central server where the Choreography software is used to compute timecourses of behavioral parameters and actions including speed (blue: 400 MB per session), from which per-animal health scores such as maximum speed in a time window are computed using the Metrology software (teal: 1 MB per session). Finally, this is collated into timecourses for visualization by averaging across animals at each timepoint (green: 10 kB per session). All categories of data, save the full video feeds which are processed live, are backed up to a cluster filesystem which is accessible to the web server that runs the user interface (gold). **(B)** Portion of a raw image as captured by the Observatory. Image covers 3.4×4.68 mm. **(C)** Example of a worm outline and path, extracted from (B) with the MWT and visualized with Choreography. Outline corresponding to image is shown; paths of animals’ centroids are plotted for all times. Paths are colored according to movement speed: yellow is faster, red is slower. Scale as in (B). **(D)** The speed of the centered animal in (C) over the “aroused speed” window (post-stimulation). Measurement window indicated by gray box. Thin black line indicates maximum speed measured from this animal (after 5-point median filter). **(E)** Maximum speed for animal of interest (green dot) and others on the plate (open black circles) as measured by Metrology.

The data that feeds into this pipeline is transferred from the image computer’s local drive to the server’s networked drive. This is accomplished by a program called Saver, written in Scala, that moves the data and compresses the image snapshots as PNG files instead of uncompressed TIFF (which is used by Spanner for speed). Saver and Spanner alternate runs: Saver can complete its tasks in less time than it takes to load a new tray, and this way the imaging computer can devote all of its resources to Spanner when a tray is present.

This data is then processed through four subsequent stages in a sequential batch mode. For robustness, we designed these stages to depend only on the saved state on the various drives, not on any other record of previous work. Although this has been a good choice initially, in the future this may need to change as it involves inspecting the entire directory structure each time a stage is run, which does not scale well as data size continues to increase. One solution is simply to archive old data. However, to allow browsing and reanalysis of old data, this would be inadequate; better would be to mark older data as handled and avoid examining it unless we had reason to believe it changed.

Each stage is time-limited, as it is important that the first stage run frequently to keep the comparatively small data transfer drive free. When everything is running normally (no interruptions or manual intervention requiring extensive recomputation), all stages complete in less time than a recording from a single tray (roughly 10 minutes). Out of an abundance of caution, we wait one cycle for newly produced data to be run through the next stage of processing (in case of long-running external processes that do not complete before the next stage starts). Therefore, the final stages of analysis are available approximately 40 minutes after the data is taken. This could, with appropriately careful engineering, be sped up to a just-in-time scheme that should complete in minutes; however, given that animals are only examined once every six hours and the experiments unfold over many days, we have not in practice found that the 40 minute delay impedes us in any significant way.

Each processing stage is a stand-alone program (albeit using shared libraries) written in Scala. Coordination of the different stages is achieved with a bash shell script running in a loop.

The first stage, Tier1, gathers the data from the six separate subdirectories shared to the imaging computers, moves it to a large local drive for further processing, and also compresses the contour data and saves the contours and images to the cluster filesystem for backup. This completes the transformation, started on the imaging computer, of raw images (Fig. 3B) to outlines and positions ready for further processing (Fig. 3C).

The second stage, Tier2, extracts low-level parameters from the contour and centroid data, using the Choreography program that is part of the original Multi-Worm Tracker distribution; Choreography now has its own repository (https://github.com/Ichoran/choreography). In brief, we use the standard command-line parameters `-N all –shadowless -S -t 30 -M 1.5 -p 0.04 -s 0.2,` the plugin commands `–plugin mwt.plugins.Reoutline::exp –plugin mwt.plugins.Respine –plugin mwt.plugins.SpinesForward,` and the output directive `-o area,speed,midline,loc_x,loc_y.` Together, these gather per-animal output of length, area, x and y position, and speed measured over an 0.2 s time window while rejecting likely low-quality data from putative animals that have moved less than 1.5 body lengths or were followed for less than 30 seconds. When computations are completed, the results are also compressed and saved to the cluster filesystem. This results in a transformation of the data into per-animal parameters, such as speed, measured over time (Fig. 3D).

The third stage, Tier3, selects animals that were followed successfully during three windows of interest: “initial”, 10–20 s after protocol start; “calm”, 275–295 s after protocol start, where animals have had time to return to a calm state after the initial taps; and “aroused”, 440–450 s, 30 seconds after the last of the twelve taps given with a 10 second interstimulus interval—in addition to inducing per-tap habituation, this protocol results in a prolonged increase in activity in response to the repeated agitating stimuli. Key parameters (length, size, and speed) are calculated over this window (mean length, mean size, and both mean and maximum speed; though the speed is first passed through a median filter of window size 5 to reduce outliers). Finally, the results are again compressed and saved to the cluster filesystem as backup. This results in a transformation of the data to per-animal scores that can be used as a quantification of animal health (Fig. 3E).

The final stage, Tier4, was intended to group this per-animal data by biological condition and compute and draw key plots for examination by the user. However, we found that this posed an architectural problem, as users would want to update information on the user interface—to, for example, remove from consideration a plate that was observed to be contaminated with mold—and see it immediately reflected in the output. Because this was incompatible with the batch-mode processing, and we observed the Tier4 computations to be very fast, we moved the logic from Tier4 into the user interface code. In the future we plan to partially restore Tier4 by keeping track of which metadata was used to create each graph, and modularize the code such that either Tier4 or the UI code can call the same module to create graphs. However, for now, Tier4 exists mostly in conception; technically, the last stage of computation is done upon request by the user, and thus the Tier4 code only coordinates metadata between the cluster and server. In any case, this conceptual tier results in a transformation of the data to graphs of parameters, with animals averaged across plates with the same biological condition during the same 6 h recording cycle forming each data point.

#### 3.2.9 Webservatory

Previous studies utilizing the Multi-Worm Tracker have required considerable post-capture processing before any biological insight could be gleaned ([Ohyama et al., 2013, Podshivalova et al., 2017]). However, since the output of the *C. elegans* Observatory has a natural biologically interpretable form—graphing of parameters against age, with different strains or conditions compared to each other—we were able to create a web interface that allows the experimenter to view these key graphs without performing any data processing tasks themselves.

The front page of the interface gives a graphical display of key environmental parameters and device utilization (Fig. 4A). It also allows each experimentalist to log in to view their experiments. Each experiment is given a unique 4-character identifier (one digit, three capital letters). Within an experiment, users can define their library of distinct conditions or strains (Fig. 4B) and specify both a nickname by which to refer to them, and what other conditions or strains should be plotted as a control. They then can specify which slots in which tray have which sample (Fig. 4C), along with a start and stop time and notes, if applicable. When the experiment begins, the *C. elegans* Observatory gathers snapshots of each plate for each session (6 hours between sessions), which can be viewed to check for mold or other problems (Fig. 4D). If a plate is found to have a problem, its entry in the tray can be removed, or the duration of validity can be shortened, so that final results do not contain bad data.

**Figure 4:**
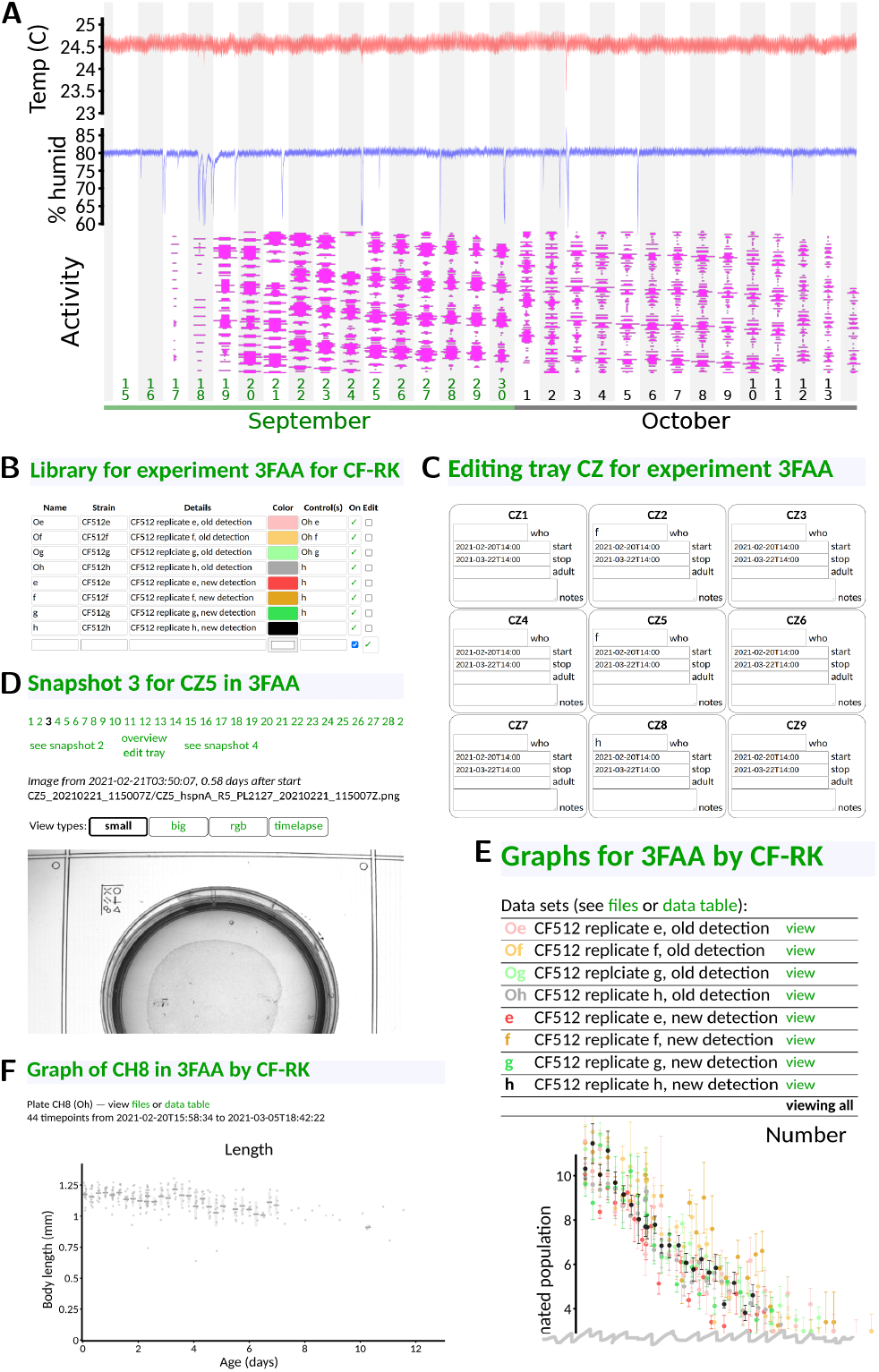
Web-based user interface. All panels taken directly from interface with no editing save cropping. **(A)** Front page status monitoring. Users can see at a glance temperature (red), humidity (blue), and instrument utilization (magenta; width indicates fraction each tray filled with plates, and time throughout the day runs from low to high). Downward spikes in humidity are caused by opening the incubator to, for instance, load samples. **(B)** Library definitions. Users can give nicknames and colors to their samples, in addition to specifying strains and condition information. **(C)** Assignment of samples to tray positions. Only the sample nickname is required, and only in plates that exist; dates are auto-populated from experiment-level metadata. **(D)** Visual checking of snapshots. Users may see snapshots of any recorded plate in order to, for instance, check for mold or drying. **(E)** Viewing of results. By default, all samples are shown. Data points are averaged across six-hour time bins (one complete cycle through all trays). A variety of graphs are presented (only the top of the first graph, sample number, is shown). **(F)** Viewing of per-plate detail. Data underlying each sample can be viewed, if desired. Gray dots indicate measured length of individual animals for one particular plate of one sample type. Additional graphs are situated below the first (not shown).

This then allows results to be grouped by strain or condition, and presented to the user graphically (Fig. 4E). We show only a small portion of the interface here: each strain/condition is listed under its user-specified color, and by default all data is plotted together (the top portion of the number-of-animals graph is shown, as that appears first on the interface). By clicking on a strain/condition name, one can get a direct comparison of condition and its control(s) (not shown), and from there one can click on individual plates to see the underlying animal-by-animal data (Fig. 4F).

In this way, an experimentalist can rapidly understand phenotypes as they become visible, and can navigate the data to check for quality issues.

The web interface is written in Scala, using the open source HTTP framework Cask (which is fashioned after the better-known Python framework Flask).

#### 3.2.10 Statistics

Whenever possible, we have attempted to visually display contrasting conditions in a way that lends itself to an intuitive understanding of both variability within conditions and differences across them. For this we either plot the underlying data points, or a Monte Carlo sampling of underlying data that serves as a null-hypothesis distribution. We typically utilize a one-sequence implementation of the RXS M XS 64 variant of the PCG64 pseudorandom number generator ([O’Neill, 2014]) for our random number source; an initial 64 bit seed is arbitrarily chosen by mashing fingers on the number keys.

Internally, the software uses basic statistical functionality and distributions, to, for instance, calculate standard error estimates. These are mostly programmed from first principles and available in KSE (https://github.com/Ichoran/kse), a Scala library adding the general-purpose functionality not found in Scala but commonly used by one of the authors. The PCG64 random number generator is also in this library. This basic functionality was adequate for the error analysis in Fig. 9.

The Kaplan-Meier estimates of survival in Fig. 8B were computed in R version 3.5 using the `survival` and `rms` packages.

We expect that reasoning based on visual impression will typically be at least as valid as that based upon a quoted p-value, but provide p-values for key contrasts nonetheless. p-values for day 3 / day 0 ratios (Fig. 7) were computed using the Mann-Whitney test (`wilcox.test` in R 3.5 or `MannWhitneyTest` in Mathematica 12.1). p-values for different day 3 and day 0 scores than control (Fig. 10, Suppl. Table 1) were computed by first projecting onto the first principle component of the points in this space, then using the Mann-Whitney test on those values. Proportions of animals detected between day 9 and day 0 were tested using the chi-square test (R 3.5) on the number of animals in each case. All p-values are reported without correction for number of comparisons of different samples. Because not all sources of variability are adequately understood—true in many studies—these p-values should be interpreted primarily as a statement about the the numerical structure of the data. While we have done our best to ensure that systematic variation is due solely to the condition we intended to vary, it would be wise, as always, to maintain appropriate skepticism regarding batch effects and other potential confounds.

### 3.3 Experiment Design

#### 3.3.1 Strains and Sample Preparation

One of the key challenges to overcome in lifespan experiments and whole-life experiments in *C. elegans* is the appearance of progeny. Since *C. elegans* is a self-fertile hermaphrodite, unless active measures are taken to prevent this, one will get a new generation of animals every three or four days (precise timing dependent on temperature, strain, and/or perturbation). One common way to prevent progeny from accumulating is to induce sterility chemically. For example, 5’-fluorodeoxyuridine (FUdR) is a nucleoside analog which interferes with development sufficiently to prevent eggs from hatching ([Gandhi et al., 1980]). However, although we have successfully used FUdR in Observatory experiments, we note that the drug must be added at the correct developmental stage to be effective at preventing progeny while avoiding defects in the adults; this is challenging when interventions or genetic backgrounds could also affect the rate of development. Another approach is to use microfluidic devices that retain adult animals but allow young animals to be washed away ([Rahman et al., 2020]). This, however, at a minimum would require automated attachment and detachment of a fluid handling system, which we judged to be impractical. A third approach, which we took, is to perform experiments in a temperature-sensitive sterile genetic background. In our case, we used *rrf-3* ; *fem1* (strain CF512) animals, which have largely normal egg-laying at 15 °C but no progeny (< 1*/*10000 from our observations) at 25 °C. The primary downside of this approach is that mutations have to be crossed or re-engineered into this background; but, as we were mostly interested in performing RNA interference experiments, this was an acceptable tradeoff. Fortuitously, the *rrf-3* mutation enhances the effectiveness of RNAi ([Simmer et al., 2002]) in addition to preventing reproduction at higher temperatures (for which the *rrf-3* gene was independently isolated as *fer-15*) . All data in this paper is from CF512 animals (i.e. *rrf-3* ; *fem-1*) . Animals were housed at 15°C unless otherwise indicated.

To prepare worms for recording, we used one of two methods to achieve an age-synchronized population. For experiments with a small number of plates per strain and relaxed requirements for synchronization, we picked two day-one adults to a seeded plate and left them for 10–12 hours to lay eggs, then removed the adults. For experiments with larger numbers of animals or where we wished to have tighter synchronization, we used an L1 larval arrest protocol (streamlined from [Stiernagle, 2006]). In brief, we left hundreds of adult animals to lay eggs on seeded 6 cm plates, then washed off the adults and any already-hatched progeny with 2 mL filter-sterilized S-basal plus 0.01% PEG. The eggs, which mostly remain behind with washing, were scooped off, cleaned with 30 s immersion in 1/2x worm bleach solution (1:2:7 ratio of 1 M KOH, bleach, and water), rinsed 4x with S-basal + PEG, and left to hatch overnight at 25 °C in S-basal + PEG. Without food, the hatched L1s arrest their development, increasing the synchrony between recently-hatched and less-recently hatched animals. The hatched L1s were then pipetted onto plates (volume determined by the density of animals in the liquid, aiming for 40–60 animals per plate) and maintained at 25 °C. These two methods are diagrammed in Fig. 5A.

**Figure 5:**
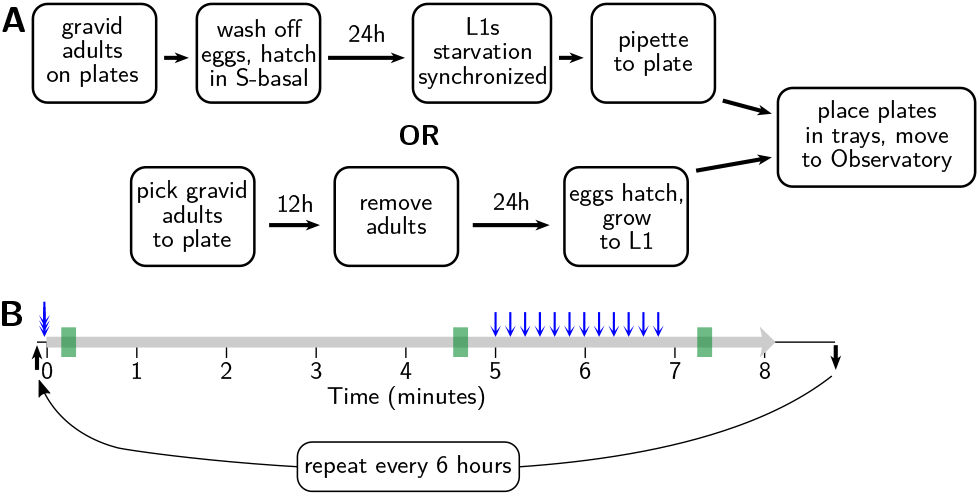
Sample preparation and behavioral assay protocols. **(A)** Age synchronization. One of two standard methods is used to synchronize progeny at 15 °C, as shown, before moving to the Observatory at 25 °C. **(B)** Behavioral assay. Tray arrives at imaging platform (black arrow) and tray is triple-tapped (blue tripleheaded arrow) prior to the onset of recording (thick gray line). After 300 seconds, 12 taps are delivered (blue arrows) with a 10 second inter-stimulus interval; after 80 more seconds recording ceases. Tray is removed shortly thereafter (black arrow) until the next session 6 hours later. Summary parameters are gathered in 10 second windows at 10 s, 275 s, and 440 s (green rectangles), corresponding to initially active, calmed, and tap-aroused behavioral states.

Assays run on OP50 had plates (NGM) and bacteria prepared using standard methods. Generally, 50 or 100 mL of OP50 culture was used, and spread near to but not all the way to the edges of the plate to provide a large area for animals to roam without visual occlusion or loss of focus caused by the edges of the plate. Plates were left to grow for 2–3 days, though neither the precise timing nor the amount of bacteria appears to have a substantial impact on the aging trajectory (preliminary results; see Figure S1).

RNAi assays had bacteria prepared using a protocol we have previously found successful (based largely on [Kamath et al., 2001]). Frozen stocks of RNAi were grown overnight at 35 °C on LB plates with 100 µg/mL carbenicillin. Single colonies or a streak were picked from plates to LB liquid media with 12.5 µg/mL tetracycline and 100 µg/mL carbenicillin and grown overnight at 37 °C. The media then was diluted 1:1 into LB with carbenicillin only and grown for two more hours at 37 °C. Next, 1 mM IPTG was added and the culture was placed at 30 °C for 4 hours. Finally, 100 µL culture was pipetted to each assay plate. Plates were NGM with 1 mM IPTG and 100 µg/mL carbenicillin; the culture was spread with a glass rod to cover most of the plate, and plates were left to grow for 24 hours at 30 °C.

After plates were loaded with animals, the lids were coated with anti-fogging agent (FogTech DX wipes). They were then parafilmed, and 2–4 small holes were punched in the parafilm between the lid and plate to allow a small degree of venting. This prevents any fogging while also reducing drying sufficiently to allow month-long experiments.

#### 3.3.2 Standard Behavioral Assay

We used a behavioral assay slightly modified from [Podshivalova et al., 2017]. Previously, recordings made using the MWT would happen after the normal lid of the plate was swapped for a custom glass-covered lid, providing a strong multi-sensory experience (changes in temperature, humidity, oxygen, etc.) that agitated the animals. However, with the *C. elegans* Observatory, no such agitation is an inherent part of the process: the delivery to the imaging platform is quite gentle. Because the MWT requires animals to move in order to detect and quantify their behavior, we needed an initial pre-recording stimulus in order to assist detection. Blue light stimulation has been reported to be robust ([Churgin et al., 2017]), but the logistics of deploying it were complicated. Because we already wished to give mechanical stimuli, we tested whether a closely-spaced trio of taps could serve as an effective agitating stimulus before recording. We found that at least for young and middleaged animals, it could. In contrast to the lid-change stimulus, after the mechanical stimulus, animals rapidly returned to baseline activity. We therefore devised the protocol as shown in Figure 5B: the agitating triple-tap, 5 minutes of cooldown, a tap habituation assay consisting of 12 taps with a ten-second inter-stimulus interval, and finally an 80 second cooldown. Note that we do not presently score the habituation to tap, but as we plan to in the future, and it also induces long-lasting elevated movement that we do quantify, we have included it in the protocol.

Given that switching trays takes about one minute, the protocol allows for slightly over 9 minutes between sessions, which with 32 trays works out to just under 6 hours per cycle, or four sessions for each tray per day. Although it is possible to detect reproducible changes on a 6 hour timescale, such changes are very small, and a slower cycle time would almost surely be adequate (every 8 hours or every 12).

### 3.4 Lifespan Scoring

The MWT behavioral analysis algorithms rely critically on motion for segmentation of animals from background. However, animals exhibit an extended period of limited mobility before death. This leaves the system unaware of both dead and very slow-moving animals, and therefore prevents the number of detected animals from being used to calculate the number of living animals. However, because existing automated systems score lifespan based on static images ([Stroustrup et al., 2013]), we expected that we would be able to score lifespan from the handful of images stored each session. Although we intend to automate the process, initially we sought only to score the images manually, in order to verify that lifespans in the *C. elegans* Observatory were as expected. To highlight changes over time, we took images from three successive sessions (spanning 12 hours) and encoded them, intensity-inverted, in red, green, and blue channels. Thus, truly motionless worms appear gray, while slightly moving worms have patches of color about them. This considerably accelerates the task of manually scoring the time of death.

## 4 Results

### 4.1 Validation

#### 4.1.1 Quantification of Behavior

Our first task upon completing the construction of the *C. elegans* Observatory was to decide which behavioral metrics to compute and display for users. Initially, we favored those that were highly robust and easy to interpret biologically. Since size and movement speed met these criteria, we focused on validating these. We intend to add additional metrics in the future, including, for instance, propensity for dwelling vs. forward or backward motion, rate and magnitude of response to individual taps, and frequency of turning. Because the MWT only detects moving animals, and does not try to maintain animal identity when animals collide, it cannot be used to provide an accurate count of animals. Firstly, some animals may never move enough to be detected; secondly, those that are detected may appear as several independent records punctuated by collisions, with no indication that these records in fact belong to the same animal. However, it is highly desirable for an experimentalist to have at least an estimate of the number of animals on the plate to tell if the plate has been loaded properly, and to tell when the behavioral data from an experiment is effectively over. We use a simple greedy heuristic to try to join up records by judging whether an animal could have traveled from the position of loss to the newly found position; this provides a more accurate estimate of the number of moving animals. We observed that young adults are most easily detected; as animals age, an increasing fraction of them fail to move enough to meet the detection threshold, and thus fewer and fewer are detectable (Fig. 6A).

**Figure 6:**
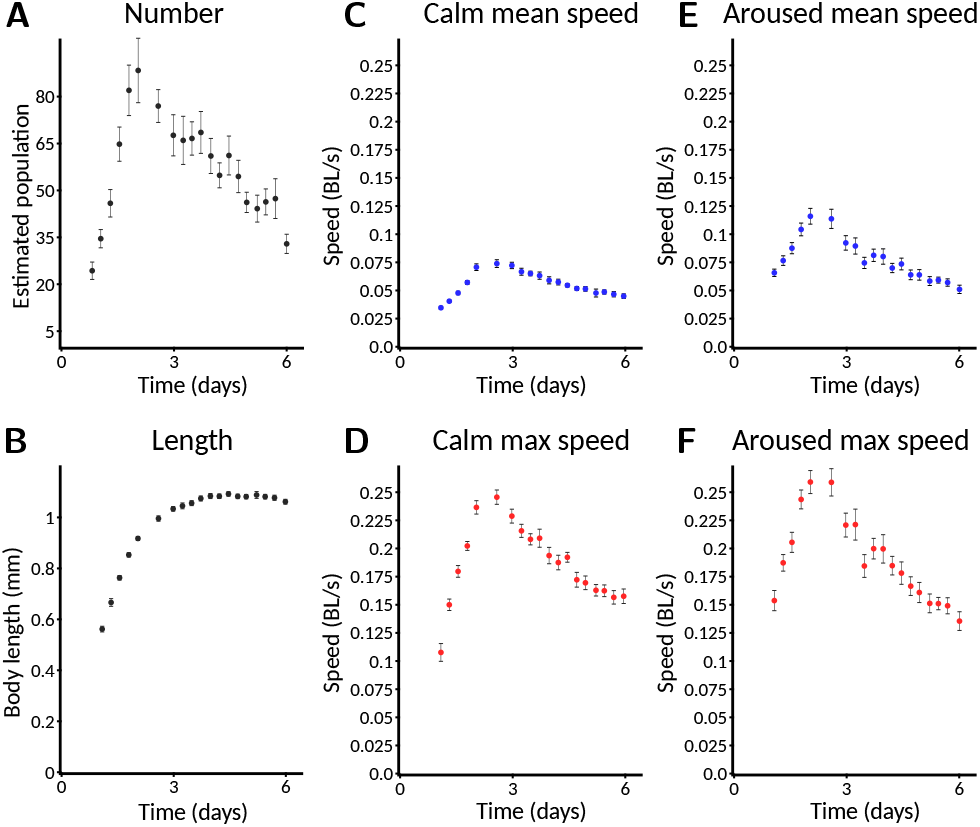
Selected standard behavioral measures. Plates imaged from L1 are used as an example, and are followed for only six days. Error bars indicate S.E.M.. Data from a plate is only included for plotting if there are at least three animals detected on that plate. **(A)** Estimated number of animals. Animals are counted only if they move enough for their behavior to be quantified. Specific speed scores may have a smaller sample size as not all animals are followed at all times. Error bars represent S.E.M. across plates, not within-plate error. **(B)** Length of animals. Note that data does not appear at the earliest timepoints because the animals are too small to detect. **(C)** Calm mean speed. Mean speed of animals in measurement window shortly before taps are delivered (275–285 s), when animals have had time to return to baseline activity after the taps at the start of recording. **(D)** Calm maximum speed. Maximum speed of animals in (C). Each animal’s maximum speed is computed within the window; from these per-animal maximum scores, a population mean and standard error are computed. **(E)** Aroused mean speed. Mean speed of animals in measurement window shortly after taps are delivered (440–450 s), when animals still exhibit an overall arousal in behavior following repeated stimuli. **(F)** Aroused maximum speed. Maximum speed of animals in (E) using the method described in (D).

To increase the ability to judge quantitative differences between conditions, all graphs save animal number produced by the Observatory include only data from plates with three or more detected animals. Thus, the absence of plotted data at a particular timepoint indicates not that zero animals have been detected, but that too few have to trust a quantitative comparison.

For size metrics, we compute both length (length of an 11-point segmented line that runs down the center of the outline contour) and area (corresponding to number of segmented pixels); these, obviously, are highly correlated. Since area adds little, we typically focus on length (Fig. 6B), which shows an increase through mid-adulthood, followed by decline during aging, as expected ([Hulme et al., 2010]).

For speed metrics, we plot speeds calculated over short windows at three different times. The first window, “initial”, is 10–20 s after the beginning of the recording and captures the behavior of worms freshly agitated by the triple-tap protocol. However, only worms near their peak activity can be detected so quickly, so this metric is of limited use in reporting aging phenotypes. The second window, “calm”, is 275–295 s after the beginning of the recording, by which time speeds have returned to baseline (Fig. 6C and D). The third window, “aroused”, is 440–450 s after the beginning of the recording and 30 s after the end of the tap protocol. This protocol induces a sizable increase in the animals’ movement speed (Fig. 6E and F; compare to “calm” speeds). For each time window, we computed speed two ways: the mean speed across the window (Fig. 6C and E), and the maximum speed (after noise reduction with median filter; Fig. 6D and F). Though the calm and aroused speeds are both similarly robust, we favor the latter as a better measure of the animals’ capacity rather than their motivation. Similarly, between mean and maximum speed, we favor maximum as we judge that more likely to represent the animals’ capacity—this has been shown to be an important consideration for *daf-2* mutants ([Hahm et al., 2015]), for instance.

Hereafter, we focus on “aroused maximum speed” as a single behavioral metric that captures how an animal’s capacity for motility changes throughout life. In cases where we wish to compare the number of animals across conditions, we compare the number of animals successfully measured for aroused maximum speed. Graphs are plotted with time measured relative to initiation of recording (typically animals are L1, though this can vary).

Because quantification of behavior relies on a minimum amount of movement for worms to be distinguished from background, the behavioral parameters cannot be computed late in life when animals’ movement drops below this minimum. However, the degree to which animals have become undetectable is itself determined by movement. Thus, in addition to comparing aroused maximum speed earlier in life, we also compare changes in the number of animals detected and measured for a later-life estimate of behavioral vigor.

#### 4.1.2 Validation of Known Mutants

We expected that the *C. elegans* Observatory would reveal behavioral phenotypes for genes known to be involved in longevity. In particular, we wished to find both an example of progeria, where young adult behavior was normal but the decline in behavior was faster than normal, and an example of extended health, where young adult behavior was normal but the decline in behavior during aging was slower than normal.

As part of running controls for other experiments, we found that *hsf-1(RNAi)* produced a particularly clear progeria phenotype (Fig. 7A), consistent with its shortened lifespan and accelerated tissue damage ([Garigan et al., 2002]). We also found normal growth but a markedly early and severe age-related decrease in length (Fig. 7B). To provide a more quantitative metric for progeria, we plotted animals’ peak speed (averaged over the plate) against the speed three days later (Fig. 7C), and found that even plate-by-plate, *hsf-1(RNAi)* showed a sufficiently large early decrease in speed to lie almost completely outside of the control distribution. Thus, we anticipate that significant progeria could be detected in a screen that uses only a single plate per condition.

**Figure 7:**
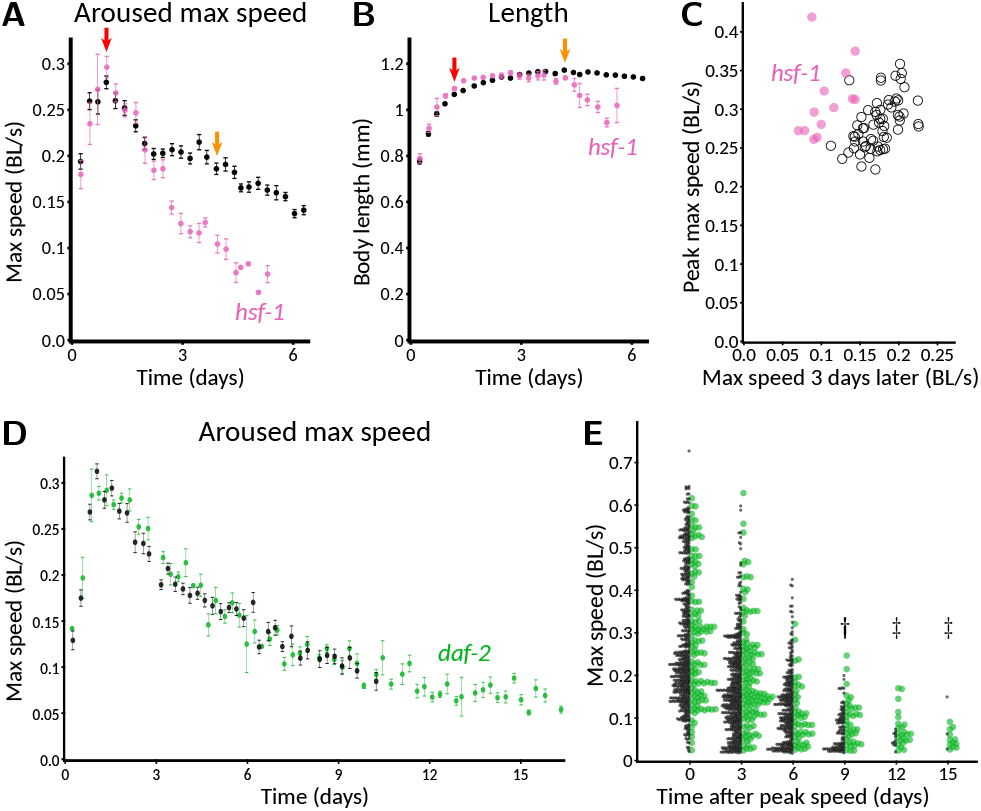
Validation of genetic perturbations resulting in reduced and extended maintenance of youthful behavior. **(A)** Aroused maximum speed of control (black, n = 14 plates) or *hsf-1(RNAi)* (pink, n = 5 plates). Red arrow indicates time of peak activity. Orange arrow indicates 72 hours later. Note the marked decrease in speed of *hsf-1(RNAi)* animals at the timepoint indidicated in orange, but not at red. 40–60 animals per plate. **(B)** Length of CF512 animals on control (black) or *hsf-1* RNAi bacteria (pink). Arrows and sample size as in (A). **(C)** Consistency of progeric phenotype in *hsf-1(RNAi)*. Each plate is characterized by peak speed (y-axis) and speed after three days (x-axis), corresponding to the values of graphs at the red and orange arrows in (B). Replicates across four experiments and two experimenters are shown. Pink dots, *hsf-1(RNAi)* (n = 13 plates). Black circles, control (n = 62 plates). *p <* 10^−6^ that the two sets of points have the same ratio of day 3 over day 0, by Mann-Whitney U test. **(D)** Aroused maximum speed of control (black, n = 12 plates) or *daf2(RNAi)* (green, n = 4 plates). Data points are plotted only when at least one plate has at least three animals detected and measured, so the extended detection of *daf-2(RNAi)* reflects greater motion overall if not higher speed among detected animals. 60 animals per plate. **(E)** Extended maintenance of moving fraction of *daf-2(RNAi)* with age. The aroused maximum speed of every detected animal is shown in a one-sided beeswarm-style plot for control (black) and *daf-2(RNAi)* (green) at three day intervals. Each dot corresponds to the score for one animal on that day. Because the *daf-2* data set (n = 123 measurements on day 0) is smaller than the control (n = 536 on day 0), dot size is normalized such that total area of dots is equal at day 0, providing a visually faithful representation of the decreasing fraction of detectable animals. Underlying data is the same as in (D). †, *p <* 0.01 and ‡, *p <* 0.0001, probability by chi-squared test that the reduction from day 0 in number of animals measured is the same for wild-type and *daf-2(RNAi)*.

Although our controls for other experiments included gene knockdowns expected to extend lifespan, we did not find any that markedly increased motility three days after peak speed. This included *daf-2(RNAi)* animals, though not entirely unexpectedly. In previous, manual experiments with the MWT, we also found that *daf-2(RNAi)* was insufficient to show an early motility phenotype. Specifically, although long-lived *daf-2(e1368)* hypothesized ligandbinding-domain mutants exhibited increased motility relative to wild type from mid-adulthood onward, *daf-2(RNAi)* animals differed from control only in that they continued to move (detectably) after the control animals had stopped ([Podshivalova et al., 2017], Fig. 1B and Suppl. Fig. S2C). The reason for the difference was unclear; one possible explanation could involve the fact that neurons tend to be insensitive to RNAi because they do not express the dsRNA transporter SID-1 ([Calixto et al., 2010]). In the *C. elegans* Observatory, *daf-2(RNAi)* animals appeared indistinguishable from control throughout the time when speeds could be compared using automatically generated graphs (Fig. 7D).

To gain a clearer picture of how detectability varied, especially given that automatic graphing rejects as potentially unreliable data from plates with two or fewer worms, we also plotted the data animal-by-animal at various timepoints. Consistent with [Podshivalova et al., 2017], we observed that a significantly higher fraction of *daf-2(RNAi)* animals moved enough to be detected after control animals did not (Fig. 7E, day 9 after peak speed and later). We also used animal-by-animal plotting to verify that *hsf-1(RNAi)* animals could not be detected for as long as controls (Fig. S2; significant difference on day 3 after peak speed and thereafter).

#### 4.1.3 Lifespan in the Observatory

To verify that lifespan was not affected by conditions in the *C. elegans* Observatory—for instance, by the repeated behavioral assays—we compared a manual lifespan conducted in a separate 25 °C incubator with one scored off of images from the *C. elegans* Observatory using our red-green-blue method that helps call attention to time of death (Fig. 8A). We observed good agreement between the two (Fig. 8B; additional data not shown), indicating that, as expected, worms were living normal-length lives within the *C. elegans* Observatory.

**Figure 8:**
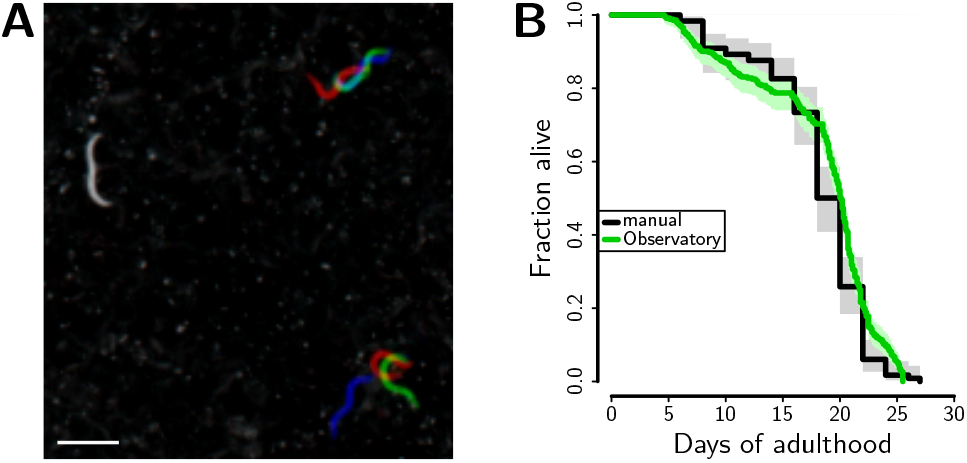
Lifespan determination. **(A)** Snapshots saved during successive recording sessions (6 hours apart) were overlaid in red, green and blue. This allows rapid visual identification of animals who are dead (gray; left), are moving their whole bodies (red, green, and blue; lower right), or moving part of their bodies (part gray, part color; upper right). Scale bar is 1 mm. **(B)** Lifespan of CF512 on RNAi control bacteria either in the Observatory (green, median lifespan 20.1 days, n=343 deaths) or in a 25 °C incubator (black, median lifespan 20.0 days, n = 119 deaths).

#### 4.1.4 Variability in Behavioral Parameters

To understand the reproducibility we could expect from the *C. elegans* Observatory, we collected the control data from three experimenters across ten separate experiments. The different experiments had different goals, but in all cases the control conditions were nominally the same. This revealed broad agreement between maximum-speed aging profiles computed for each of the plates, but nonetheless, there was substantial variability around the mean (Fig. 9A). In order to understand the source of this variability, we decomposed it into variability across experiments (Fig. 9B) and variability within experiments (i.e. between a plate and the population mean for all plates in that experiment, Fig. 9C). Surprisingly, we found that the variability from each, by eye, was roughly comparable. We therefore quantified the variance explained both by the experiment and by the plate within the experiment, at each timepoint, and plotted it as coefficient of variation (standard deviation over mean; Fig. 9D). This revealed that, indeed, the two do contribute comparable variability.

**Figure 9:**
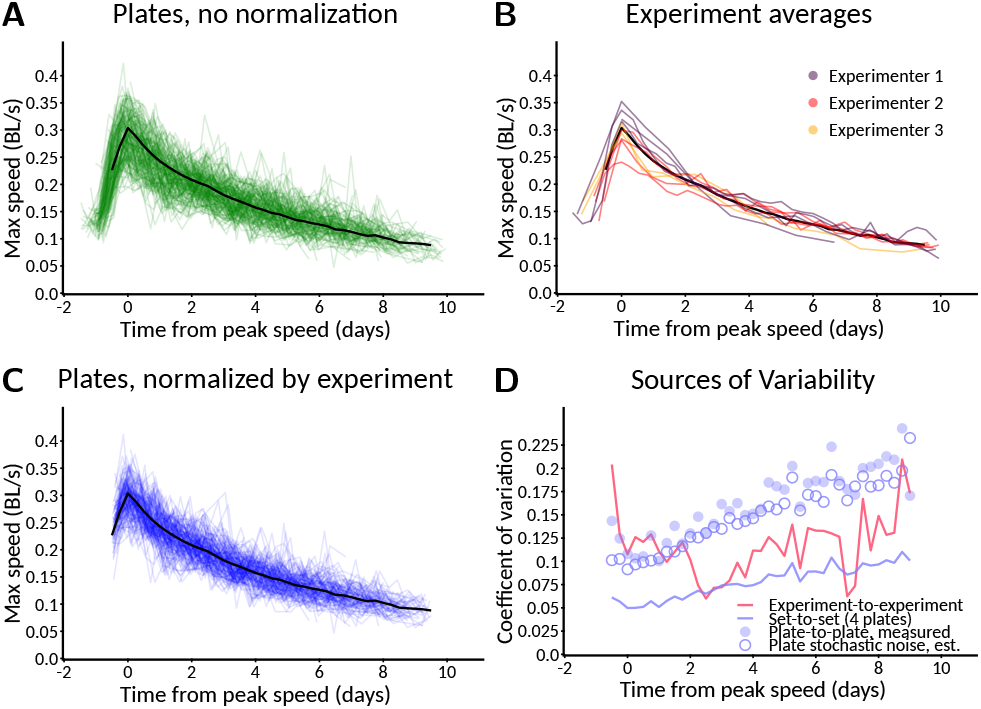
Estimation of sources of variability. **(A)** Measured variation in aroused maximum speed. Scores for 201 individual plates of control animals (green) across 10 experiments, plus population average (black). Typically 40–60 animals per plate. **(B)** Variation between experiments. Experiment averages (colors) and population average (black) for data in (A). **(C)** Variation normalized by experiment. Scores for 154 plates normalized relative to their experiment (blue) by scaling mean of experiment to match population average (black). Data from (A), with low-n plates filtered out. **(D)** Relative scale of sources of variability. Variation between experiments (red line) is large compared to variation between a 4-plate set of samples (blue line). Measured plate-to-plate variability (filled blue circles) closely matches expectation from stochastic sampling (open blue circles), indicating minimal systematic plate-to-plate effects. Variability is expressed as coefficient of variation compared to population average (black lines in A–C).

Variation from plate to plate within an experiment could result from either systematic plate-level factors, or simply from the stochastic nature of sampling (both in the random selection of animals that appear on a plate, if animals have characteristic variations; and in the sampling of random fluctuation of behavior during the assay). By sampling from the distribution of observed speeds, normalized to the plate mean, we created fictitious plates where by construction there was no systematic plate-level variation and only stochastic sampling remained. The predicted variability of these stochastic-only plates closely matched the observed variability of real plates (Fig. 9D, closed blue circles vs. open blue circles), indicating that under these conditions, a majority of the variability comes from stochastic sampling; differing conditions between the plates does not seem to be a significant concern. Note, however, that we prepare all plates at the same time, and remove from consideration any plates contaminated with mold or undesired strains of bacteria, precisely to avoid systematic differences between plates.

#### 4.1.5 Power Analysis

Our goal, setting out, was in part to create a system that could in principle scale up to the level of performing wholegenome screens. In our existing experiments, we have typically run four plates per condition, which gives us a coefficient of variation of slightly over 5% in maximum speed parameter (predicted value: Fig. 9D, green line). With this level of noise, what size of effects can we reasonably expect to recover? First, consider the approximation where all measured values, and ratios thereof, have a Normal distribution. If we choose a significance cutoff of 0.05 (not corrected for multiple comparisons) as our threshold for detection, and desire an 80% chance of detecting a variation that increases only the mean and not the noise (another simplifying approximation), this is equivalent to requiring that only 0.2 of the distribution of the shifted mean falls below the 0.95+ point on the tail of the null hypothesis distribution. Thus, for a one-sided test we require a difference of

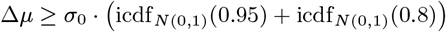

where icdf represents the inverse cumulative distribution of a probability distribution, *N* (0, 1) is the Normal distribution with mean 0 and standard deviation 1, and *σ*_0_ is the standard deviation of the measurement of the peak-to-day-three ratio of activities. This works out to roughly 2.5× the coefficient of variation for a single-sided test or 2.8× a two-sided test to reach *p <* 0.05 80% of the time. For a ratio of peak speed to day 3 speed (with measured coefficients of variation of 0.055 and 0.077 respectively), and naive propagation of error to the ratio, this works out to an 80% detection chance for a roughly 24% change in mean for a one-sided test, or 27% for two-sided.

Monte Carlo sampling of the wild-type data in Fig. 9, assuming a control set of 30 plates, and a sample size of 4 plates per experimental condition, with true mean peak speed assumed not to vary across conditions and day three varying multiplicatively, gives an empirical 80% chance of passing a two-sided test threshold of *p <* 0.05 when the value is changed by 1.31×, in line with the approximate result.

These results inform our expectations about what reasonably could be found in a whole-genome screen. Smaller changes could be robustly detected by using more of the data, thereby lowering the measurement noise. For instance, instead of comparing a single timepoint, values could be averaged over a wider temporal window (perhaps three sessions—18 hours—centered on peak speed, and five sessions centered on day three).

### 4.2 Novel Aging Trajectory Phenotypes

Although the *C. elegans* Observatory has the potential throughput to enable an unbiased screen, we were curious whether a candidate-gene approach could reveal novel behavioral trajectory phenotypes. Via computational analysis of human health data (Libert et al., unpublished, preprint doi https://doi.org/10.1101/2022.04.14.488358) and literature search, a variety of candidate mammalian genes were selected; we picked *C. elegans* genes corresponding to some of these, based on existence of orthologs, paralogs, or at least reasonably closely related gene families, and the presence of corresponding RNAi constructs in the Ahringer RNAi library ([Kamath et al., 2003]).

Despite having only ten candidate genes (Supplementary Table 1), we found several interesting RNAi phenotypes. For early aging, we found both considerably elevated and reduced maintenance of youthful movement (Fig. 10A) in *tank-1(RNAi)* and *cdc-42(RNAi)* respectively. For later-life aging, we found no candidates that clearly increased detectability after 9 days, but we found three that markedly reduced it: *cdc-42(RNAi), scav-1(RNAi)*, and *scav-2(RNAi)*.

**Figure 10:**
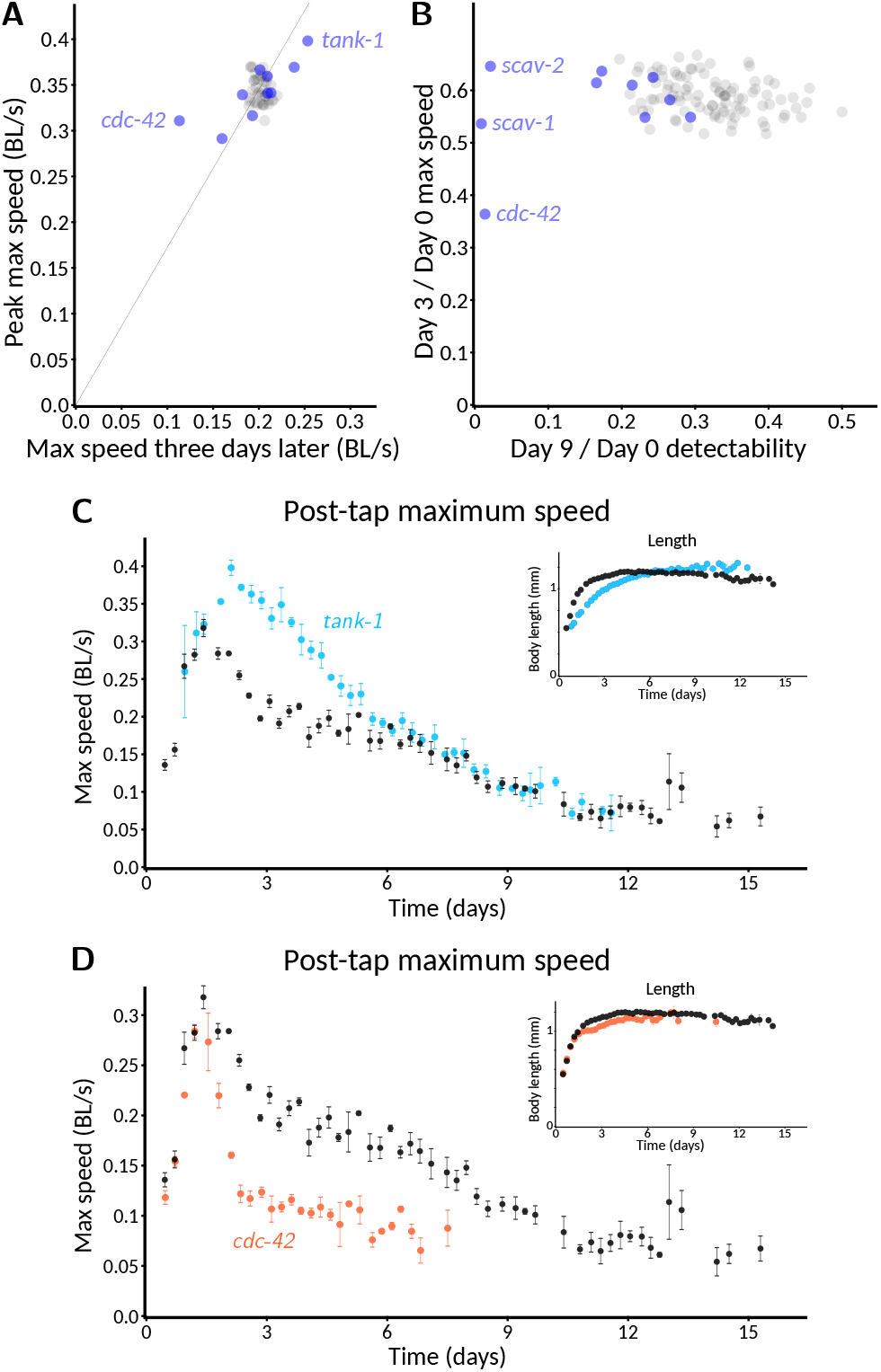
Pilot screen of gene candidates implicated in healthy mammalian aging. **(A)** Effect on early vigor. Aroused maximum speed at peak vs. three days later for RNAi of ten candidate genes with 3–4 plates per condition and 30–60 worms per plate (blue) compared to random 3–4 plate samples out of the 16 control plates representing the expected distribution (gray). Thin line indicates constant peak to three-day speed ratio. Slowest and fastest three-day conditions are named (slow: *cdc-42*, fast: *tank-1* ; *p <* 0.01 that each is from the control distribution) corresponding to best candidates for reduced and extended vigor, respectively. RNAi began at L1. **(B)** Effect on detectable motility. Ratio of animals detected on day nine vs. day zero compared to peak speed (horizontal axis) against ratio of early vigor (day three / day zero speed), with samples from (A). The three genes showing dramatic loss of detectable motility are named (*cdc-42, scav-1*, and *scav-2*). Experiment averages (blue) and random control sets (gray) for data in (A). **(C)** Speed timecourse for *tank-1(RNAi)* (average of four plates). Inset shows length of animal, illustrating slower development of *tank-1(RNAi)* animals. **(D)** Speed timecourse for *cdc-42(RNAi)* (average of three plates). Inset shows length, illustrating normal developmental timecourse.

The full behavioral trajectory of *tank-1(RNAi)* revealed an intriguing phenotype: animals grew slowly (Fig. 10C inset), yet they did not simply “live slow”, with similarly slowed behavior: the young *tank-1(RNAi)* animals were if anything more vigorous than their control counterparts. They then maintained their youthful vigor for an extended period, after which their activity returned to control levels. In contrast, *cdc-42(RNAi)* showed a consistent progeric phenotype reminiscent of *hsf-1(RNAi)*: despite growth and peak speed that closely matched controls, movement speed dropped more rapidly in treated animals than in controls (Fig. 10D).

More generally, these results demonstrate the effectiveness of the *C. elegans* Observatory at discovering when a gene can influence aging trajectories.

## 5 Discussion

The *C. elegans* Observatory was conceived of as a highthroughput yet approachable tool for studying factors that influence how animals age. Thus far we have focused on usability and throughput rather than a diverse set of behavioral metrics, reasoning that the ability to easily measure one primary behavior—stimulus-aroused motility—throughout life was the most important advance in our capabilities. Since the MWT captures both position and postural information (exemplified in Fig. 3C), we plan to re-analyze existing data to quantify additional behaviors. The MWT has been used to quantify a variety of behaviors both in young animals and aging ones ([Swierczek et al., 2011, Podshivalova et al., 2017]), but the algorithms make assumptions about magnification and stability that are not fully met in the *C. elegans* Observatory configuration, so each behavior requires validation and some will require algorithmic adjustments. We anticipate that the additional richness will provide deeper insight to the biology of aging. For instance, we would like to ask whether genes that affect youthful motility coordinately affect the youthful presentation of other behaviors. Being able to acquire the data, as we are now, is the first and most critical step.

Thanks to its use of standard plates and its relatively straightforward user interface, it is easy to use for smallscale experiments, and this is mostly how we have used it thus far. Nonetheless, its ability to scale to larger screens is promising. A single individual can load the entire *C. elegans* Observatory in roughly two days of intensive work, with four plates per RNAi condition. Although this would be a grueling activity to keep up for months on end, if one had five Observatories, a single individual could in principle run over 2,500 conditions (10,000 experimental plates) per month for two-week assays. This would allow the entire updated Ahringer RNAi library, containing roughly 20,000 samples, to be screened in approximately eight months, with 80% power to detect changes of 30% at a 5% false positive rate. Alternatively, a less sensitive single-plate assay, which would require no more than twice the effect size for the same detection chance, could be performed on a single Observatory in well under a year. These capabilities position the *C. elegans* Observatory favorably for a thorough understanding of the genetics of behavioral aging.

Among worm trackers, the *C. elegans* Observatory occupies a useful position along the outer envelope of throughput and precision, reminiscent of a quote from journalist A. J. Liebling: “I write faster than anyone who can write better, and I write better than anyone who can write faster”. In comparison to our system, the behavioral detail provided by a multi-camera 96-well format processed with the Tierpsy tracker ([Barlow et al., 2022]) is considerably greater (due both to focus on detailed analysis of morphology and behavior, and also the 8 µm pixel size as opposed to 40 µm here); and has the advantage of easily switching to longitudinal mode (one worm per well; they typically use three). However, the *C. elegans* Observatory will run five times the number of conditions with 3–4 × the number of animals per condition, as compared to their full 30-camera system. Interestingly, a hybrid system is conceptually straightforward: their arrangement of five 96-well plates fits comfortably within the area of a single Observatory tray. On the other hand, the WormCamp (Fouad et al., unpublished, preprint doi https://doi.org/10.1101/2021.10.18.464905) system makes tradeoffs to go faster: they use a stage-mounted camera assembly to traverse over a large field of 24-well plates, giving them access to roughly 4x more wells than the Observatory holds plates but with reduced assay time (5 minutes as compared to 9 in the Observatory) and activity only scored at the population level, not animal-by-animal. For our purposes, the *C. elegans* Observatory strikes the right balance between throughput and power to detect behavioral phenotypes, but other systems that push the outer edge of the throughput-precision space also have considerable potential to expand our understanding of the aging process.

As yet, we do not know what fraction of genes will produce an appreciable phenotype when knocked down with RNAi. Our very limited candidate gene approach was surprisingly successful, though it is difficult to tell to what extent this is a function of the prevalence of good targets and to what extent it is due to astute selection. Although a longevity phenotype had not, to our knowledge, previously been demonstrated in *C. elegans, cdc-42* was predicted to be a candidate aging/longevity pathway gene ([Witten and Bonchev, 2007]) in addition to its known role in establishment of cell polarity ([Cowan and Hyman, 2007]). Likewise, no *C. elegans* longevity phenotype has been reported for the PARP-family gene *tank-1* ; in mammals, tankyrase was originally identified as a telomere-associated protein ([Azarm and Smith, 2020]) and has a diversity of other roles ([Damale et al., 2020]). In *Drosophila*, mutations in tankyrase have been reported to reduce both lifespan and climbing behavior ([Li et al., 2018]), but we are not aware of any reports that reduced tankyrase might also increase the duration of youthful vigor. Understanding the molecular basis of the *cdc-42(RNAi)* and *tank-1(RNAi)* phenotypes would obviously require additional study, starting with a longer recording or a manual assay to determine the lifespans. Nonetheless, finding that at least two out of ten candidate genes had interesting RNAi-knockdown phenotypes was encouraging.

Seemingly intractable biological problems often require the development of new techniques and technologies before they at last yield and we begin to gain insight. It is our hope that the *C. elegans* Observatory, and tools like it, will play a part in developing a mechanistic understanding of the basis of aging.

## Author Contributions

RK and CK conceived of and designed the study. RK designed, built, and programmed the automated system. AR, JG, and RK conducted experiments. RK wrote the manuscript and constructed the figures. All authors reviewed, improved, and approve of the manuscript.

## Acknowledgments

We are grateful to many members of Calico Life Sciences LLC. Particularly important contributions include the following: Alfred Millet-Sikking and Andrew York assisted with mechanical and optical design of the system, and Alfred additionally helped with electronics. Eddie Xue, Jacob Kimmel, and Adam Baker provided feedback on software engineering and user interface design. Peter Noone and Alex Chekholko supported our needs for computational infrastructure including networking and cluster storage. Katie Podshivalova and Peichuan Zhang ran trial experiments as the system was being developed, providing feedback on how to improve the system. Sergiy Libert provided a list of candidate mammalian genes. Ashok Shah ensured ample supplies of media and worm plates of various sorts. Beyond Calico, Zachary Pincus gave advice on prioritization and scope of project. Strains from the *C. elegans* Genetics Center (CGC), which is funded by the NIH Office of Research Infrastructure Programs (P40 OD010440), were used during testing and development.

## Data Availability

The full video streams are transient, and even the reduced forms (e.g. outlines) are impractical to share. The data extracted from Tier3 analysis that was used to create the figures can be found in https://doi.org/10.5281/zenodo. 6645842 along with the code we used to produce figures from the relevant portion of the data. Software and CAD files for the entire Observatory system are also available there, in the form used in this publication. Additionally, further refinements to the Observatory, along with assembly diagrams and documentation when they are created, will either be available at or linked from https://github.com/calico/ elegans-observatory.

