## Supplemental Methods, Figures, and Table for "The *C. elegans* Observatory: High-throughput exploration of behavioral aging"

### Supplementary Material for "The *C. elegans* Observatory: High-throughput exploration of behavioral aging"

June 15, 2022

#### Supplementary Methods

##### Octicons

One key problem we had to solve was how to identify which tray was being imaged at any given time, given only what was visible through a single camera (Suppl. Fig. 3). We investigated traditional barcoding schemes to achieve this, but found that cutting barcodes directly onto acrylic was difficult to perform robustly, and were worried that any sort of adhesive label risked becoming detatched and therefore compromising reliability. Furthermore, we found that barcode-reading software is generally not designed around responses within a few milliseconds, and therefore the barcode reading could introduce delay beyond that set by the camera's frame rate.

We then considered naming the positions in a human-readable way only, using OCR to extract the identity. Again, however, we found that easily available OCR software was not designed around fast responses. Therefore, we elected instead to devise a custom solution for automatic identification of trays. In particular, to identify each tray we chose a 36-character set of capital and lower-case letters that we judged to have low possibility for confusion even when handwritten (Suppl. Fig. 3B). Slots within each tray were numbered 1–9, right-to-left from top, thus making the identity visually clear and easy to write. We then designed a whimsical 8-character set, dubbed "octicons", of easily laser-engraved icons (Suppl. Fig. 3C) with low overlap when superimposed (Suppl. Fig. 3D) for rapid template-matching. Six of these characters, five encoding tray plus slot within tray and one reserved for a checksum, were engraved in a small rectangular area edged on two sides by straight lines to allow easy alignment. This allows sufficiently rapid and robust identification that we can use the identity-code reading process to also detect when a tray is in position for imaging (when ready, the code is not only successfully read but also the position of the code does not change appreciably from frame to frame).

Laser cutters, in addition to cutting through materials, typically have a feature to either vector engrave by cutting shallow trenches in the material, or to raster engrave/etch images by burning the surface enough to create contrast. Of the two, we found vector engraving to produce higher contrast under our brightfield illumination scheme, and to be faster create and less susceptible to physical damage. Therefore, we chose line-based symbols suitable for laser vector engraving (Suppl. Fig. 3C) and which have low overlap when symbols are superimposed (Suppl. Fig. 3D). Eight symbols seemed like a reasonable compromise between the limits of our creativity in coming up with different symbols sans some formal optimization process, and the desire to encode more data per symbol (three bits are encoded). Since we decided to label the trays with a pair of letters for human identification and selected 36 easily discriminable letters to use, we could potentially create 1296 distinct trays; but we also need to discriminate each position on the tray, for a total of 11644 possible targets. Thus, we needed at least 14 bits of information to encode every possible valid tray identity and position, or at least five octicon symbols.

Because the magnification is fixed, a tray's rotation is precisely controlled as it is placed for imaging, and a tray's position is fairly highly controlled, we were able to simply produce a binary template image corresponding to the ideal version of each octicon and then calculate the dot product between the template and the actual image at the target position.

In order to precisely define the target position for each symbol, we placed the symbols in a $3 \times 2$ rectangle (5 characters or 15 bits encoding, plus 1 character or 3 bits checksum) framed by a line along the left and lower edges of the rectangle. This line is trivial to detect by summing rows or columns of a portion of the image and looking for a dip in intensity. In order to know where to perform this flattening operation, we individually measured the target area for every camera and recorded the position in a settings file. Then, since the distance from the edges to each symbol is fixed, the appropriate subimage can be selected. Since there can still be deviations of a pixel or two for various reasons (an incompletely flat field of the image, deviations in laser cutter performance, or slight differences in focus), we perform a local search by shifting the template one pixel away in each direction, and then repeating if we find a higher dot product. Even though we did not take all possible steps to optimize this process—for instance, subdividing the dot product into the white part (which is easily shifted a row or column at a time) and the dark part (separately for each octicon, which requires addressing only a small subset of pixels) would greatly speed the computation—the resulting detection was both fast and accurate.

The online detection code for octicons is part of Spanner and was written in C++, though a Scala routine is used to generate the templates used in the C++ code. Both are available, along with SVG files of the ideal octicons and black-and-white images of octicons as expected at the magnification used in the system, in the main archive.

The octicon strategy could be adopted for other similar identification processes. If this were done, we recommend at least one change: the backwards L-like symbol can, when multiple symbols align, also produce a fairly deep valley in intensity when summing images along rows or columns, thus requiring some care when using the rectangular frame to define position. It would be even more robust to, for instance, rotate the symbol slightly. It would be better yet to create families of discriminable engravable symbol sets of different size with maximal difference under expected operations (inversion, small translations, maybe rotation), optimized via simulated annealing or genetic algorithms.

### Supplementary Figures

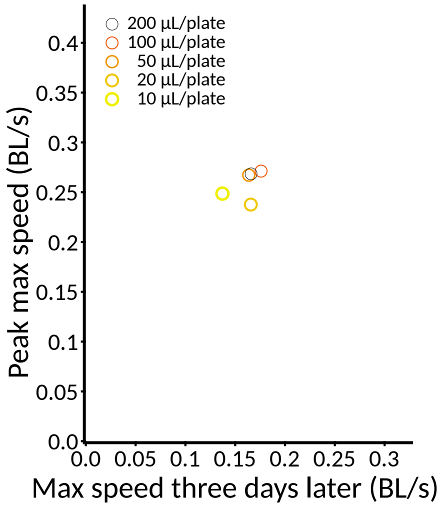


Figure 1: Preliminary comparison of behavior on plates seeded with varying levels of bacteria. Mean peak speed is plotted against mean speed three days after peak for CF512 animals in different conditions. Plates were seeded with 10 µL (n = 2 plates), 20 µL (n = 4 plates), 50 µL (n = 5 plates), 100 µL (n = 14 plates), and 200 µL (n = 4 plates) of OP50 and grown for two days at room temperature before adults were placed to lay eggs for 12 hours and placed in the Observatory for monitoring thereafter. Note that a typical experiment would have 50–200 µL of bacteria per plate, with the same volume used for all plates used in one experiment. Lawn thickness was visually different in all five conditions, but the amount of food present per plate was not quantified.

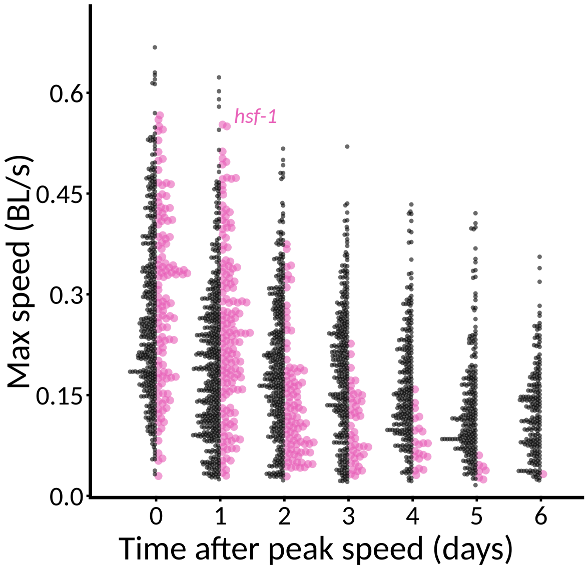


Figure 2: Distribution of aroused maximum speeds in *hsf-1(RNAi)* animals compared to control. Dual beeswarm-style plot in the style of Fig. 7E, where each dot represents one measurement from one animal on one day, except contrasting control (n = 380 animals on day 0) with *hsf-1(RNAi)* (n = 120 animals on day 0) instead of *daf-2(RNAi)* with its control. Reduction in number of detectable *hsf-1(RNAi)* animals is significant from day 3 after peak onwards ($p < 10^{-4}$ by chi-squared test). Note also that at later timepoints the distribution of *hsf-1(RNAi)* appears narrower than control (compare also to Fig. 7E).

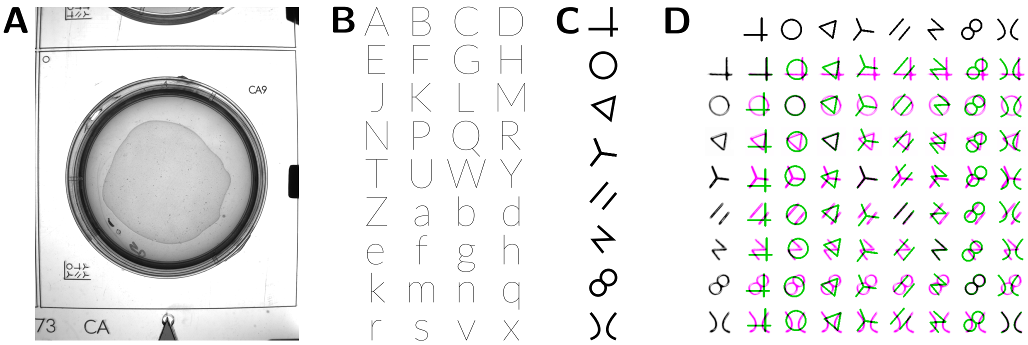


Figure 3: Tray identification. **(A)** Portion of tray visible from one camera. Note human-readable tray identifier (far bottom left), human-readable slot identifier (top right), and machine-readable code (bottom left) representing ten times the tray value plus 1-9 depending on camera position. Note bounding frame of machine-readable code for rapid alignment. **(B)** Discriminable base-36 letters. Trays are redundantly labeled with a base-36 letter code, imprinted into acrylic using laser engraving of the edge of a very narrow font (Lato, font weight Thin), as shown here. Value increases left-to-right, then top to bottom: A=0, B=1, etc.; the corresponding one-indexed decimal number is also engraved on the tray to reduce human error. **(C)** Octicons. Eight highly non-overlapping line-based symbols were chosen at the whimsy of the authors to create a octal code for machine reading that could be rapidly be discriminated by template matching. The top symbol represents 0; value increases downwards. **(D)** Octicon matching. Eight symbols taken from Observatory images (left edge) are matched against the idealized templates generated for the magnification of our images (top edge). Optimal overlap within allowed search parameters are shown. Black: overlap between target template and image. Green: unmatched target template. Magenta: unmatched darkness in image.

### Supplementary Tables

#### Candidate Gene Parameter Values

| Gene | Library ID | Peak Speed | Day 3 Speed | P | Detectability |
|---|---|---|---|---|---|
| control | L4440 | $0.336 \pm 0.008$ | $0.195 \pm 0.005$ | | 0.290 |
| C26D10.4 | C26D10.4 | $0.341 \pm 0.012$ | $0.213 \pm 0.012$ | | 0.243 |
| F52C6.4 | F52C6.4 | $0.341 \pm 0.025$ | $0.209 \pm 0.010$ | | $0.166^{*}$ |
| *cdc-42* | R07G3.1 | $0.311 \pm 0.015$ | $0.113 \pm 0.012$ | † | $0.014^{‡}$ |
| *clk-1* | ZC395.2 | $0.316 \pm 0.025$ | $0.193 \pm 0.013$ | | 0.214 |
| *pkn-1* | F46F6.2 | $0.367 \pm 0.023$ | $0.201 \pm 0.010$ | | 0.294 |
| *scav-1* | C03F11.3 | $0.339 \pm 0.022$ | $0.182 \pm 0.023$ | | $0.009^{‡}$ |
| *scav-2* | Y76A2B.6 | $0.369 \pm 0.023$ | $0.239 \pm 0.021$ | | $0.021^{†}$ |
| *scav-5* | R07B1.3 | $0.360 \pm 0.017$ | $0.209 \pm 0.010$ | | 0.265 |
| *tank-1* | ZK1005.1 | $0.398 \pm 0.027$ | $0.253 \pm 0.014$ | † | 0.173 |
| *wwp-1* | Y65B4B_11.a | $0.291 \pm 0.017$ | $0.160 \pm 0.009$ | | 0.232 |

**Supplementary Table 1**

Parameter values measured for control and RNAi knockdown of ten candidate genes. Speed values are all maximum speed, and reported as population mean and standard error. P is the p-value that the peak and day-3 speeds together are the same between control and each candidate, scored by Mann-Whitney U test after linearization along axis of greatest variation. Detectability is defined as the number of animals measured on day 9 divided by the number on day 0 (on the same plates); symbols indicate significantly different ratio by chi-square test. L4440 is the empty RNAi plasmid. Significance levels: $*$, $p < 0.05$; †, $p < 0.01$; ‡, $p < 0.0001$.
